# Fexofenadine inhibits TNF signaling through targeting to cytosolic phospholipase A2 and is therapeutic against autoimmune diseases

**DOI:** 10.1101/584540

**Authors:** Ronghan Liu, Yuehong Chen, Shuya Wang, Yazhou Cui, Xiangli Zhang, Zi-Ning Lei, Aubryanna Hettinghouse, Jody Liu, Wenyu Fu, Zhe-Sheng Chen, Chuanju Liu

**Affiliations:** NYU School of Medicine; St. John’s University

**Keywords:** TNF-α, cPLA2, Fexofenadine, inflammation, autoimmune diseases

## Abstract

TNF-α signaling plays a central role in the pathogenesis of various diseases, particularly autoimmune diseases. Screening of a library composed of FDA approved drugs led to the identification of Terfenadine and its active metabolite Fexofenadine as inhibitors of TNF-α signaling. Both Fexofenadine and Terfenadine inhibited TNF/NF-κB signaling *in vitro* and *in vivo*, and ameliorated disease symptoms in various autoimmune disease models, including TNF-α transgenic mice, collagen-induced arthritis, and inflammatory bowel disease. Subsequent studies identified cytosolic phospholipase A2 (cPLA2) as a novel target of Fexofenadine. Fexofenadine blocked TNF-stimulated cPLA2 activity and arachidonic acid production through binding to catalytic domain 2 of cPLA2 and inhibition of its phosphorylation on Ser-505. Further, deletion of cPLA2 abolished Fexofenadine’s anti-TNF activity. Collectively, these findings not only provide new insights into the understanding of Fexofenadine action and underlying mechanism, but also provide new therapeutic interventions for various TNF-α and cPLA2-associated pathologies and conditions, particularly autoimmune diseases.

## INTRODUCTION

Autoimmune diseases are a series of disorders and conditions caused by immune intolerance to self-antigens which attack specific target organs and display diverse clinical signs^1,2^. Over 100 classifications of autoimmune diseases have been documented, such as rheumatoid arthritis (RA)^3^ and inflammatory bowel disease (IBD, including Ulcerative Colitis and Crohn’s Disease)^4^. Autoimmune diseases are chronic diseases with complicated pathology and diverse clinical signs, underlying which are alterations in cytokine expression and immune cell infiltration. Among the pro-inflammatory cytokines involved, tumor necrosis factor alpha (TNF-α) has received great attention due to its position at the apex of the proinflammatory cytokine cascade and its dominance in the pathogenesis of various disease processes^5,6^, and in particular, autoimmune disorders^7,8^. TNF-α inhibitors (TNFI), including etanercept (Enbrel), infliximab (Remicade), and adalimumab (Humira), have been accepted as effective anti-inflammatory therapies and are among the most successful biotech pharmaceuticals^9–11^. Although treatment with TNFI is highly effective in ameliorating disease in some patients, current TNFI fail to provide effective treatment for up to 50% of patients^12,13^. In addition to high cost (upwards of $20,000 per year per patient using anti-TNF biologics), current TNFI have been found to increase cancer risk^14^ and cause infection in some patients^15^. Thus identification and characterization of novel, safer, and more cost-effective antagonists of TNF-α, in particular with different inhibitory properties, are of great importance from both a pathophysiological and a therapeutic standpoint. Considering the fact that drug development is time consuming and extremely expensive, costing ~15 years in time and 800 million USD on average^16^, we adopted a strategic approach involving the repurposed use of clinically approved drugs. A drug library composed of FDA approved drugs was screened both *in vitro* and *in vivo* by use of TNF-α/NF-κB reporter constructs and mice, which led to the identification of Terfenadine and its active metabolite Fexofenadine as inhibitors of TNF-α signaling.

Terfenadine and Fexofenadine are two well-known histamine receptor 1 antagonists and used for treating allergic diseases^17^. Terfenadine, a first generation anti-histamine drug, has been clinically suspended due to potential adverse events. In contrast, Fexofenadine, the major active metabolite of Terfenadine and a non-sedative third generation antihistamine drug^18^, does not have proarrhythmic risk associated with use of Terfenadine, and is marketed as an over-the-counter (OTC) drug due to its safety. Fexofenadine has been being widely used to treat various allergic diseases, like allergic rhinitis, conjunctivitis and chronic idiopathic urticarial^17–20^.

In our efforts to elucidate the molecular mechanisms underlying Fexofenadine- and Terfenadine-mediated inhibition of TNF-α signaling, we identified cytosolic phospholipase A2 (cPLA2) as a novel target of these two drugs. The major function of cPLA2 is to promote phospholipid hydrolysis to produce arachidonic acid (AA)^21^; AA activates NF-κB and is involved in the pathogeneses of various conditions^22^, including inflammatory and autoimmune diseases^23^.

Herein we present comprehensive evidences demonstrating that both Fexofenadine and Terfenadine act as the inhibitors of TNF/NF-κB signaling and are therapeutic against autoimmune diseases. Additionally, we also provide evidences revealing that these two drugs bound to cPLA2 and inhibited its enzymatic activity, which is required for their inhibition of TNF-α signaling.

## RESULTS

### Fexofenadine and Terfenadine act as the antagonists of TNF-α and inhibit TNF-α signaling and activity

To isolate the small molecule drugs that inhibit canonical TNF-α/NF-κB signaling pathway, a drug library containing 1046 FDA-approved drugs was initially screened using NF-κB-*bla* THP-1 cell line in which NF-κB beta-lactamase reporter gene was stably integrated. After repeating three times, twenty-four drugs that potently inhibited TNF-α/NF-κB activation of beta-lactamase were identified (**Fig. S1a-b**). These twenty-four isolates were subjected to a second round screen using RAW 264.7 macrophages transiently transfected with an NF-κB luciferase reporter gene. Under such conditions, only the most potent anti-TNF-α/NF-κB signaling drugs are positively screened. Eight drugs among the twenty-four candidates originally isolated were selected (**Fig. S2a-b**). In order to identify the drugs that retain anti-TNF-α/NF-κB activity *in vivo*, we performed a third round screen with NF-κB-Luc reporter mice. We first crossed TNF transgenic (TNF-tg) to NF-κB-Luc reporter mice to generate TNF-tg:NF-κB-Luc double mutant mice. Overexpression of TNF-α effectively activated NF-κB luciferase *in vivo*. IVIS was implemented for whole animal bioluminescence imaging following intraperitoneal injection of eight selected drugs into TNF-tg:NF-κB-Luc double mutant mice. Five drugs, including Terfenadine and its active metabolite Fexofenadine, were shown to effectively inhibit TNF-tg:NF-κB activated luciferase *in vivo* (**Fig. S3**). Among these five identified drugs, three, including one anti-cancer drug, are known to have severe side-effects and are not suitable for treating chronic inflammatory diseases, such as rheumatoid arthritis, we accordingly selected Fexofenadine and Terfenadine for further analyses (**Fig. 1a**).

**Fig. 1.**
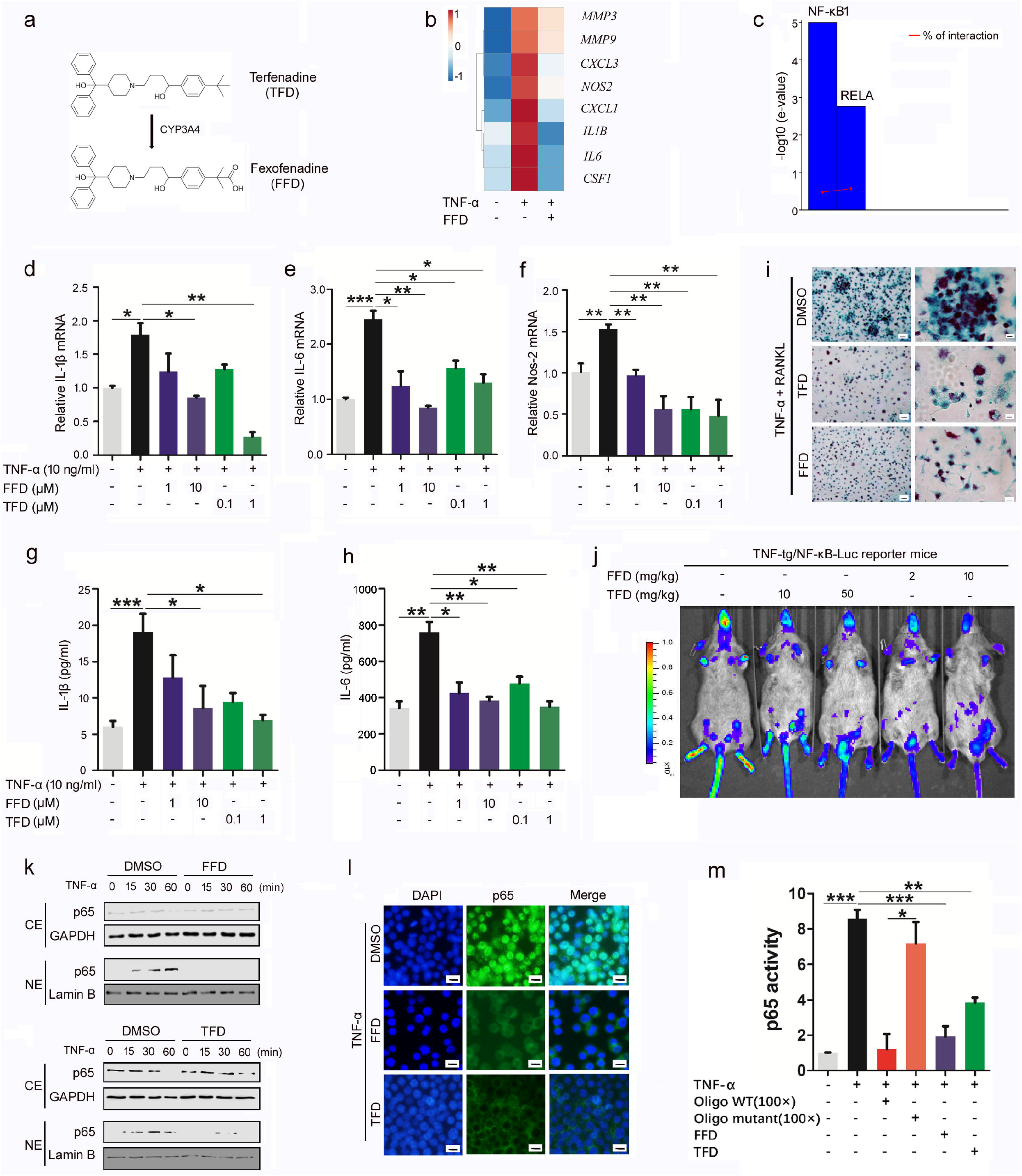
Fexofenadine and Terfenadine act as the antagonists of TNF-α and inhibit TNF-α signaling and activity. **a.** The molecular structure of Fexofenadine (FFD) and Terfenadine (TFD). CYP3A4, the major enzyme responsible for the metabolic process, is indicated. **b**. BMDMs were treated without or with (10ng/ml) in absence or presence of FFD (10 μM) for 24 hours. Total RNA was extracted for RNA-seq. A few typical TNF-α inducible genes that were suppressed by FFD were presented. **c**. Transcription factor enrichment analysis from RNA-seq results, indicating the decreased gene expressions resulted from the suppressed activity of transcription factors NF-κB1 and RELA by FFD. **d-f**. BMDMs were treated with or without (10 ng/ml) in absence or presence of FFD (1 μM, 10 μM)/TFD (0.1 μM, 1 μM) for 24 hours. mRNA expressions of IL-1β, IL-6 and Nos-2 were tested by qRT-PCR. **g-h**. BMDMs were treated without or with TNF-α (10 ng/ml) in absence or presence of FFD (1μM, 10μM)/TFD (0.1μM, 1μM) for 48 hours. The levels of IL-1β and IL-6 in supernatant were detected by ELISA. **i**. BMDMs were treated with M-CSF (10 ng/ml) for 3 days, then cultured with RANKL (100ng/ml) and TNF-α (10 ng/ml) with or without FFD (10 μM) or TFD (1 μM) for 4 days and TRAP staining was performed. Scale bar, 100μm. **j**. TNF-tg/NF-kB-Luc mice were applied to examine the anti-TNF effects of FFD/TFD in vivo. After FFD (2 or 10 mg/kg) and TFD (10 or 50 mg/kg) were orally administrated for 7 days, luciferase signals were detected by IVIS system. **k**. BMDMs were with treated with TNF-α (10 ng/ml) in the absence or presence of FFD (10 μM)/or TFD (1 μM) for various time points, as indicated. Cytoplasmic (CE) and nuclear extractions (NE) were examined by Western blot with anti-p65 antibody. **l**. BMDMs were cultured with TNF-α (10 ng/ml) in the absence or presence of FFD (10 μM) or TFD (1 μM) for 6 hours. Immunofluorescence cell staining was performed to visualize the subcellular localization of p65. DAPI was used to stain the nucleus. Scale bar, 25μm. **m**. p65 DNA binding activity was tested by ELISA. Excess amounts (100X) of WT and mutant Oligo were used as positive and negative control respectively. (* p<0.05, ** p<0.01, ***p<0.001).

We first examined the inhibition of Fexofenadine and Terfenadine on TNF-activated NF-κB pathway and downstream genes through RNA-seq with bone-marrow-derived macrophages (BMDMs) (**Fig. 1b-c, Fig. S4**). Nearly all TNF-α induced genes, especially genes encoding inflammatory cytokines, such as IL-1β, IL-6, were clearly down-regulated by Fexofenadine and Terfenadine. The lists of TNF-α inducible genes that were inhibited by Fexofenadine were used for transcription factor enrichment analysis with TFactS^24^, which led to the isolation of NF-κB1 p105 and RelA p65 as significantly regulated transcription factors by Fexofenadine (**Fig. 1c**).

In order to further validate the anti-TNF-α activity of Fexofenadine and Terfenadine, we next selected a couple of well-known TNF-α downstream inflammatory mediators for further assays. Quantitative real time PCR revealed that both Fexofenadine and Terfenadine dose-dependently inhibited TNF-α induced mRNA expressions of IL-1β, IL-6 and Nos-2 in BMDMs (**Fig. 1d-f**). Additionally, ELISA demonstrated that these two drugs abolished TNF-α induced releases of IL-1β and IL-6 in a dose-dependent manner (**Fig. 1g-h**). Similar anti-TNF activity of Fexofenadine and Terfenadine was also observed in RAW264.7 cells (**Fig. S5a-b**) and the BMDMs isolated from TNF-tg mice (**Fig. S6a-b**). TNF-α is known to enhance RANKL-stimulated osteoclastogenesis^25^. Both Fexofenadine and Terfenadine markedly inhibited TNF-α-mediated osteoclastogenesis in BMDMs (**Fig. 1i**). In addition, dose-dependent inhibition of the TNF-tg/NF-κB pathway by Fexofenadine and Terfenadine *in vivo* was also revealed by use of TNF-tg/NF-κB-Luc reporter double mutant mice (**Fig. 1j**). Furthermore, the TNF-α-induced nuclear translocation and DNA binding activity of p65 were also inhibited by Fexofenadine and Terfenadine (**Fig. 1k-m**).

### Fexofenadine and Terfenadine prevent the spontaneous development of inflammatory arthritis in TNF transgenic mice

TNF transgenic (TNF-tg) mice are known to develop an inflammatory arthritis phenotype spontaneously when mice reach 12-16 weeks old^26,27^. Next, we sought to examine the effects of applying Fexofenadine and Terfenadine to TNF-Tg mice. First, 8-week-old TNF-tg mice were treated daily with Fexofenadine, Terfenadine or Methotrexate (MTX, serving as a positive control) by oral delivery before the onset of the inflammatory arthritis phenotype. Both Fexofenadine and Terfenadine treatment resulted in reduction of all visual symptomatic signs (**Fig. 2a**) and significant reduction of clinical scores of arthritis; Fexofenadine and Terfenadine proven to be more effective than MTX, the current clinically-used small molecule drug for treating rheumatoid arthritis (**Fig. 2b-c**). In order to observe the response of inflammatory arthritis progression to Fexofenadine and Terfenadine, we stopped treatment at the 17-week-time point and resumed treatment at 19-week-time point. Ceasing the treatment led to an abrupt increase of the arthritis clinical scores. Once the treatment resumed, there was an immediate reduction in swelling score, indicating that the inflammatory arthritis induced by TNF-α overexpression responds well to both Fexofenadine and Terfenadine (**Fig. 2b-c**). H&E staining of ankle and knee tissues confirmed the inhibition of inflammatory degeneration (**Fig. 2d**). TRAP staining of paw and skull showed a preventative effect of treatment upon osteoclast differentiation (**Fig. 2e**). In addition, the drugs reduced cartilage loss, as revealed by Safranin O staining of ankle and knee (**Fig. 2f**). We also measured the serum levels of IL-1β and IL-6 and found that the levels of these inflammatory cytokines were significantly reduced in Fexofenadine- and Terfenadine-treated groups compared to the control group (**Fig. 2g-h**).

**Fig. 2.**
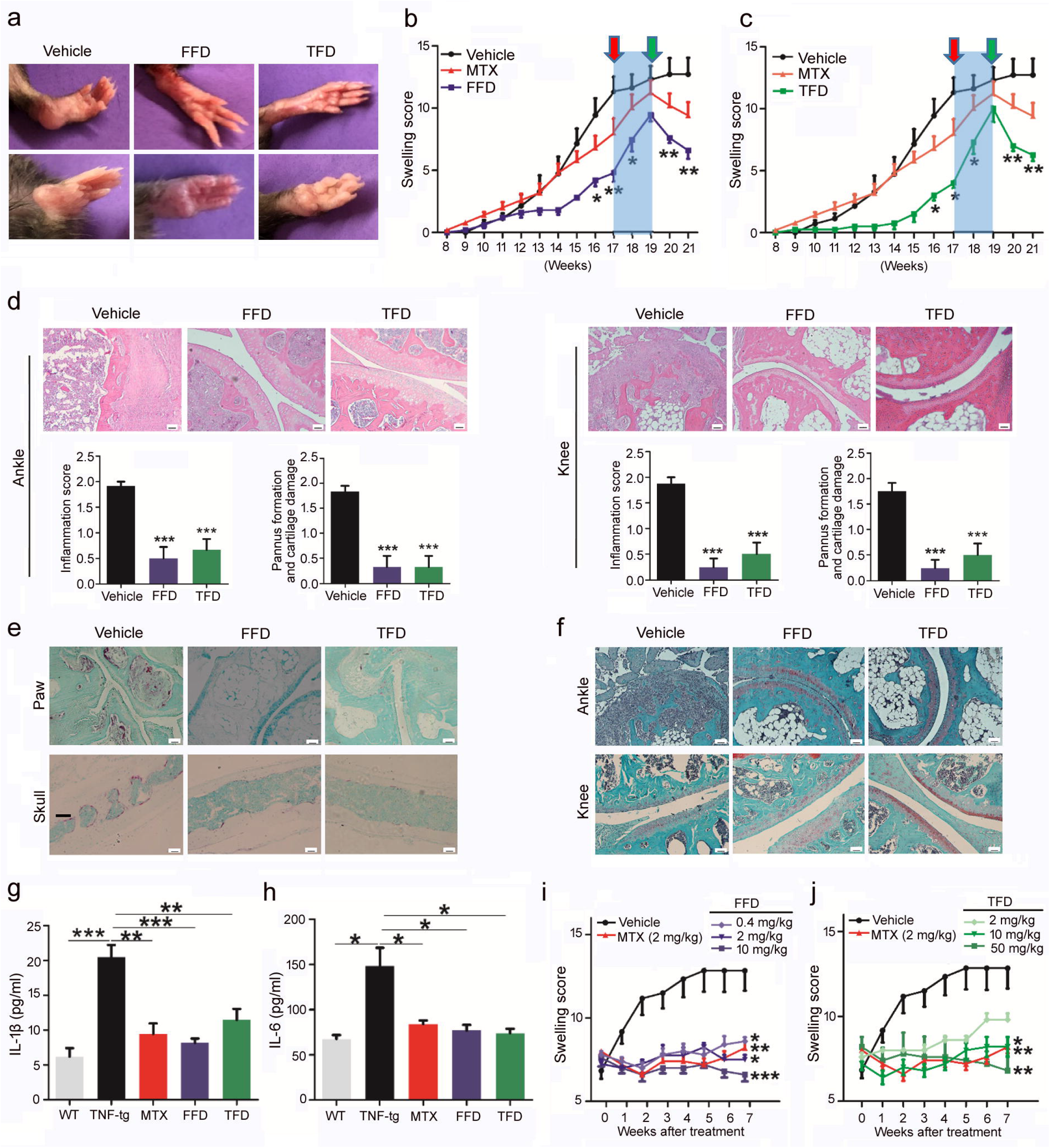
Fexofenadine and Terfenadine prevent the spontaneous development of inflammatory arthritis in TNF transgenic mice. **a-h.** TNF-tg mice (n=6) were orally administered Fexofenadine (FFD, 10 mg/kg), Terfenadine(TFD, 50 mg/kg), or Methotrexate (MTX, 2 mg/kg, serving as a positive control) daily beginning at 8-weeks of age and continuing for a total of 13 weeks. During this period, treatment was halted at 17-week point indicated by red arrow and resumed at 19-week point indicated by green arrow. **a**. Representative images of front paws and hind paws. **b-c**. Swelling score. **d**. H&E staining and quantification of histological score of knee and ankle samples. **e**. TRAP staining of paw and skull samples. **f**. Safranin O staining of knee and ankle samples. **g-h**. Serum levels of IL-1β and IL-6, assayed by ELISA. **i-j**. Therapeutic effects of FFD/TFD were tested by treating the TNF-tg mice with average swelling score reached around 8 points (n=6). Swelling scores were recorded weekly. (* p<0.05, ** p<0.01, ***p<0.001). (Scale bar, 100μm)

To determine drug’s therapeutic effects, we started treatment when the TNF-tg mice with the swelling score of approximately 8 points. Both Fexofenadine and Terfenadine showed effective therapeutic effects in a dose-dependent manner (**Fig. 2i-j**). Taken together, these data with TNF-tg mice indicate that Fexofenadine and Terfenadine exert their anti-inflammatory and therapeutic effects through the inhibition of TNF-α activity *in vivo*.

### Fexofenadine and Terfenadine prevent the onset and progression of collagen-induced arthritis

To advance understanding of the preventive and therapeutic impact of Terfenadine, especially its active metabolite Fexofenadine, on inflammatory arthritis *in vivo*, we utilized another mouse model of rheumatoid arthritis: collagen-induced arthritis (CIA), which has both immunological and pathological features with rheumatoid arthritis. CIA model was established with 8-week old male DBA/1J mice. We first started the treatment with Fexofenadine, Terfenadine, MTX or vehicle by oral delivery at 18 days after the immunization for examining their preventive effects. Severe inflammation and increased thickness in the ankles and paws were observed in the vehicle group compared to intervention groups (**Fig. 3a-b**). Analogously, Fexofenadine and Terfenadine could not only delay the onset of disease, but also significantly decrease arthritis clinical scores and incidence (**Fig. 3c-d**). Histological analysis showed less inflammation in treatment groups as compared to control group (**Fig. 3e**). Fewer osteoclasts and less bone destruction were detected in the treated groups, as revealed by TRAP staining and microCT images (**Fig. 3f-g**). Additionally, Fexofenadine and Terfenadine also prevented the loss of cartilage (**Fig. 3h**), and significantly reduced the serum levels of IL-1β and IL-6 (**Fig. 3i-j**).

**Fig. 3.**
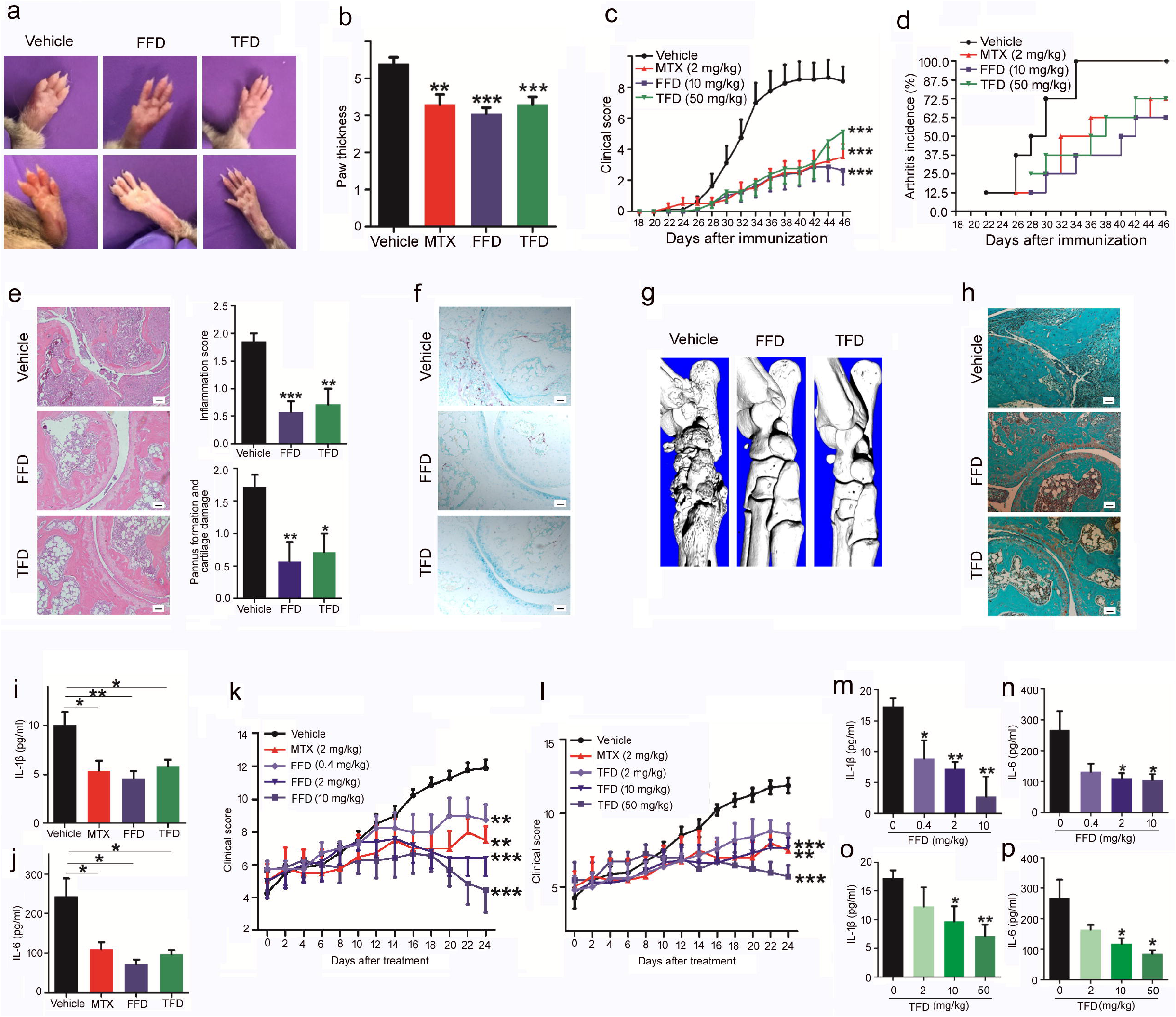
Fexofenadine and Terfenadine prevent the onset and progression of collagen-induced arthritis. **a-j.** Collagen-induced arthritis (CIA) model of DBA/1J mice was used to test prevention effects of Fexofenadine (FFD) and Terfenadine (TFD), (n=8). FFD (10 mg/kg), TFD (50 mg/kg) and MTX (2 mg/kg) were orally delivered daily beginning 18 days after immunization. **a**. The representative images of front paws and hind paws. **b**. Paw thickness. **c**. Clinical score of CIA. **d**. The incidence rate of arthritis. **e**. H&E staining and quantification of histological score of ankle samples. **f**. TRAP staining of ankle samples. **g**. microCT of ankles. **h**. Safranin O staining of ankle samples. **i-j**. The serum levels of IL-1β and IL-6 in CIA models. **k-p**. To examine the dosage-dependent therapeutic effects of FFD/TFD, CIA mice were treated with various dose of FFD or TFD, as indicated. FFD, TFD, MTX and vehicle were delivered after the clinical score reached approximately 5 points. **k**. The clinical score of FFD treated mice. **l**. The clinical score of TFD treated mice. **m-p**. The serum levels of IL-1β and IL-6. (n=8) (* p<0.05, ** p<0.01, ***p<0.001). (Scale bar, 100μm)

To determine drug’s therapeutic effects, we started treatment when the CIA model mice displayed a clinical score of approximately 5 points of a maximum 16 points per animal^28^. Both Fexofenadine and Terfenadine dose-dependently ameliorated disease scores (**Fig. 3k-l**). Meanwhile, the serum levels of inflammatory cytokines IL-1β and IL-6 were significantly decreased in the treatment groups versus vehicle (**Fig. 3m-p**). Collectively, these data indicate that Fexofenadine and Terfenadine have both preventive and therapeutic effects in CIA, an industrially used preclinical animal model for testing anti-RA drugs.

### Fexofenadine and Terfenadine are therapeutic against inflammatory bowel diseases

To determine the therapeutic effects of Fexofenadine and Terfenadine in additional autoimmune diseases other than inflammatory arthritis, we established two inflammatory bowel disease models in 8-week old C57BL/6 mice: Dextran sulfate sodium (DSS)- and 2, 4, 6-trinitrobenzene sulfonic acid (TNBS)-induced experimental colitis models^29^. First, Fexofenadine, Terfenadine, or 5-aminosalicylic acid (5-ASA, serving as a positive control) was orally delivered 3 days before model induction for a total of 11 days. Fexofenadine and Terfenadine dose-dependently prevented bleeding (**Fig. 4a and e**), body weight loss (**Fig. 4b and f**), abnormal stool (**Fig. 4c and g**) and reduced colon shortening (**Fig. 4d and h**). In particular, high dosages of Fexofenadine completely prevented bleeding, and was more effective than the positive control 5-ASA (**Fig. 4a**). Furthermore, the serum levels of IL-1β and IL-6 were reduced in treatment groups compared to the vehicle group (**Fig. 4i-l**), similar to what we observed in inflammatory arthritis models. Histological evaluation of H&E stained sections revealed the significant protective effects of Fexofenadine and Terfenadine (**Fig. 4m**). In addition, immunohistochemistry staining revealed that Fexofenadine and Terfenadine inhibited the phosphorylation of NF-κB p65 and its upstream mediator IκB-α *in vivo* (**Fig. S7a**).

**Fig. 4.**
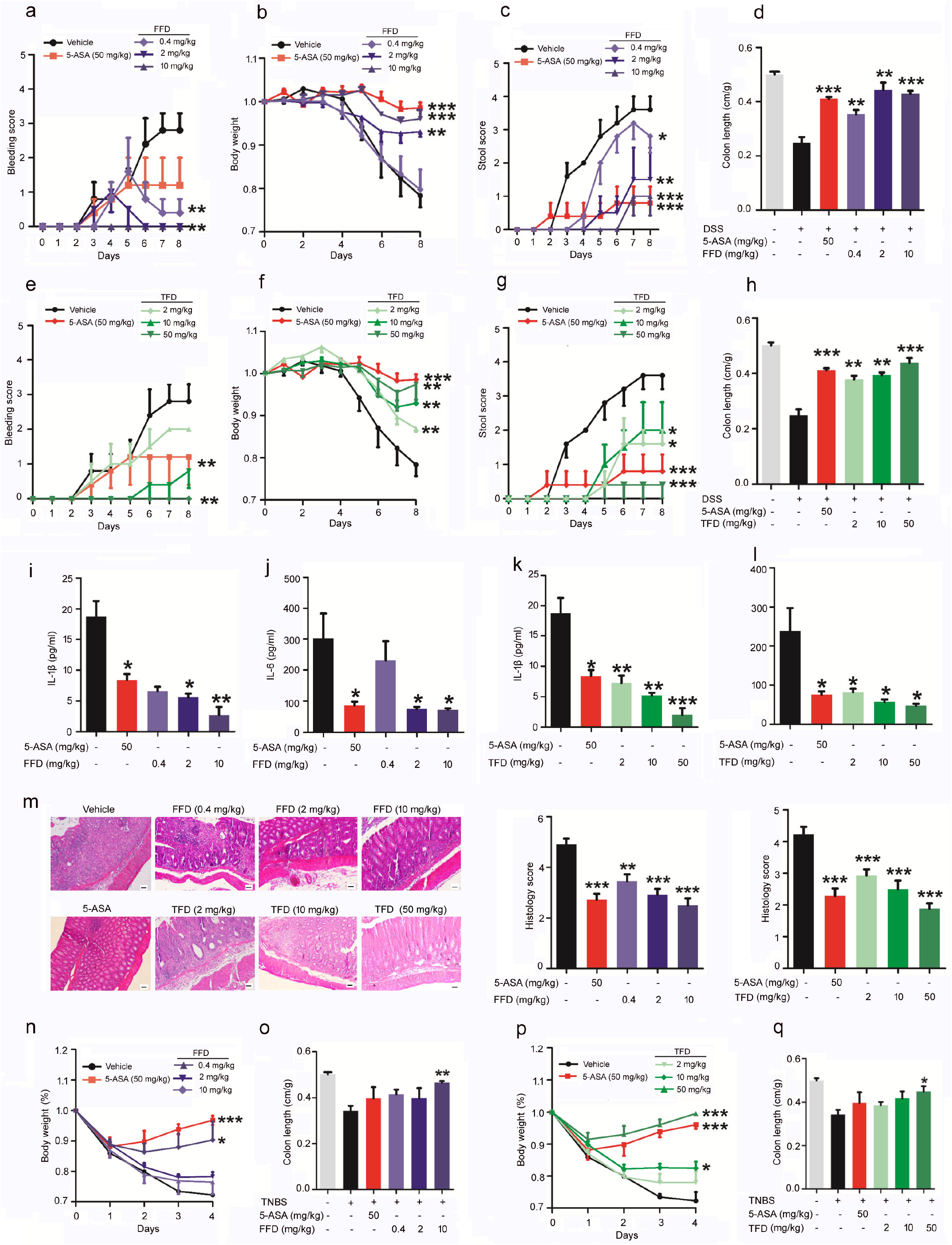
Fexofenadine and Terfenadine are therapeutic against inflammatory bowel diseases. **a-m.** DSS-induced colitis model was established in 8-week-old C57BL/6 mice (n=6 in each group) with ad libitum access to with drinking water containing 3% DSS for 5 days and followed by normal drinking water for 3 days. 5-ASA (serving as positive control, 50 mg/kg), Fexofenadine (FFD, 0.4, 2, 10 mg/kg) and Terfenadine (TFD, 2, 10, 50 mg/kg) were orally delivered beginning at 3 days before delivery of DSS in drinking water and continuing until the mice were sacrificed. **a**. Bleeding score of controls and FFD-treated groups. **b**. Body weights of controls and FFD-treated groups. **c**. Stool score of controls and FFD-treated groups. **d**. Colon lengths of controls and FFD-treated groups. **e**. Bleeding scores of controls and TFD-treated groups. **f**. Body weights of controls and TFD-treated groups. **g**. Stool scores of controls and TFD-treated groups. **h**. Colon lengths of controls and TFD-treated groups. **i-l**. Serum levels of IL-1β and IL-6. **m**. H&E staining and its quantitative analysis in colon tissue sections. **n-q**. Therapeutic effect of TFD/FFD on TNBS-induced acute colitis model. 2.5% TNBS was administered by intrarectal injection to 8-week-old C57BL/6 mice (n=6). Beginning at 3 days before intrarectal TNBS injection, drugs were orally delivered until sacrifice at 5 days. **n**. Body weights of controls and FFD-treated groups. **o**. Colon lengths of controls and FFD-treated groups. **p**. Body weights of controls and TFD-treated groups. **q**. Colon lengths of controls and TFD-treated groups. (* p<0.05, ** p<0.01, ***p<0.001). (Scale bar, 100μm)

To ascertain whether these observations were specific to the DSS-induced colitis model, another experimental colitis model which resembles Crohn’s disease was established through intrarectal administration of TNBS. The drugs were orally delivered daily and body weight was recorded. Fexofenadine and Terfenadine dose-dependently prevented both the body weight loss and the colon length shortening (**Fig. 4n-q**). Scoring of H&E stained tissue sections also illustrated dose-dependent therapeutic effects (**Fig. S8a-b and Fig. S9a-b**). Similar to results from the DSS model, immunohistochemistry staining revealed that Fexofenadine and Terfenadine inhibited the phosphorylation of NF-κB p65 and IκB-α in the TNBS model as well (**Fig. S7b**). Taken together, both DSS- and TNBS-induced colitis models demonstrated that Fexofenadine and Terfenadine have promising therapeutic effects in inflammatory bowel diseases.

### cPLA2 is a novel target of Fexofenadine and Terfenadine

The anti-histaminic activity of Fexofenadine and Terfenadine are known to be mediated by targeting to their selective histamine H1 receptor 1 (H1R1)^30–32^. We next sought to determine whether the anti-TNF activity of Fexofenadine and Terfenadine depends on their known target H1R1. We thus suppressed the expression of H1R1 using its specific siRNAs in RAW264.7 cells, and found, unexpectedly, that suppression of H1R1 did not affect the inhibition of Fexofenadine and Terfenadine on TNF-induced cytokine release (**Fig. S10a-c**). In addition, other 7 known H1R1 antagonists did not exhibit the anti-TNF activity and some of them even increased TNF-induced IL-6 release (**Fig. S11**). Collectively, these results indicate that anti-TNF activity of Fexofenadine and Terfenadine is H1R1-independent. Current clinically employed TNF inhibitors, such as etanercept (Enbrel), infliximab (Remicade), and adalimumab (Humira), exert their anti-TNF activity through disturbing the binding of TNF to its receptor TNFR1. Therefore, we next examined whether Fexofenadine and Terfenadine affected the interactions between TNF and TNFR1, leading to their anti-TNF activity. Surprisingly, both solid phase binding and flow cytometry assays showed that Fexofenadine and Terfenadine did not affect the binding of TNF-α to TNFR1 *in vitro* and to the cell surface, although anti-TNF antibody completely blocked the binding of TNF to the cell surface (**Fig. S12a-c**). These findings led us to propose that Fexofenadine and Terfenadine may have an additional new target that mediates their anti-TNF activity through a previously-unrecognized mechanism. To address this issue, we devoted significant effort to isolating protein binding partners of Fexofenadine and Terfenadine. After failure with several approaches, including labeling and biochemical co-purification, implementation of drug affinity responsive target stability (DARTS) assay^33^ proven successful. We first mixed cell lysate with Fexofenadine or Terfenadine for 1h and protease was added for 15 min. The digested proteins were separated by SDS-PAGE, followed by Silver staining (**Fig. 5a**), and a band with the molecular weight of approximately 80 kDa was found to be protected by Fexofenadine and Terfenadine. This band was excised from an accompanying Coomassie Blue stained gel for protein identification by mass spectrometry (**Fig. 5b**), which led to the identification of *PLA2G4A* encoding cPLA2 (**Fig. 5c**) and *IKBKB* encoding IKK-β as potential candidates. Both cPLA2 and IKK-β have appropriate molecular weights and are known to be the critical mediators of inflammation^23,34^. To determine whether both cPLA2 and IKK-β are the targets of Fexofenadine and Terfenadine, we performed Western blot of DARTS samples with which a series of protease to cell lysate ratios were implemented (**Fig. 5d**), and found that Fexofenadine and Terfenadine protected cPLA2, but not IKK-β, clearly indicating that cPLA2, but not IKK-β, is a novel target of Fexofenadine and Terfenadine.

**Fig. 5.**
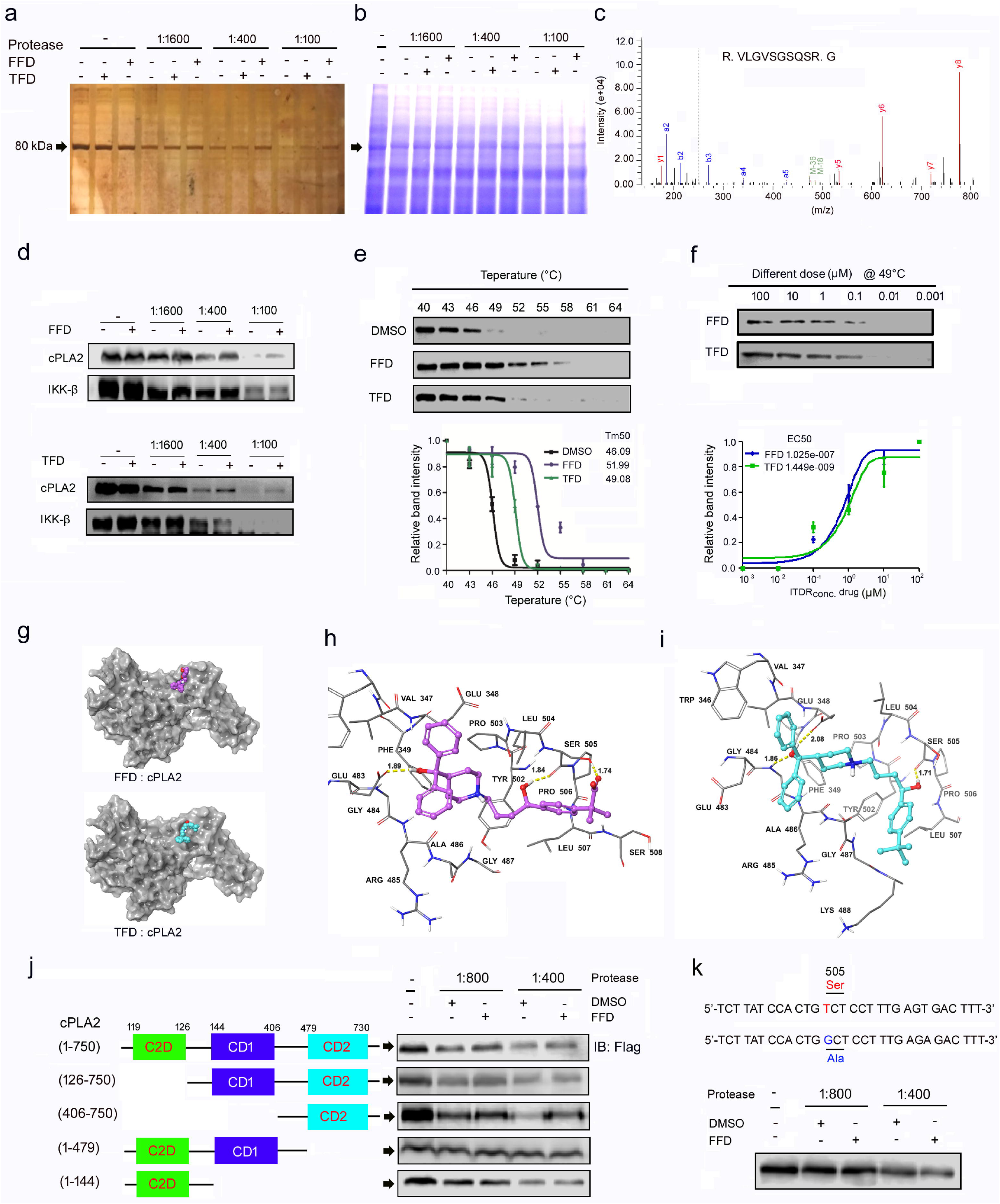
cPLA2 is a novel target of Fexofenadine and Terfenadine. **a.** Silver staining of DARTS assay. **b**. Coomassie blue staining of DARTS assay. The band with molecular weight around 80 kDa protected by Fexofenadine (FFD)/Terfenadine (TFD) was indicated by arrow. **c**. Adapted image of a mass spectra for *PLA2G4A*, encoding cPLA2. **d**. DARTS and Western blot to confirm FFD/TFD’s binding targets. **e**. CETSA melt response and associated curve. **f**. Isothermal dose response (ITDR) and its curve. **g**. IFD simulated binding complexes of cPLA2-FFD and cPLA2-TFD, respectively. cPLA2 is shown by surface representation (grey). FFD and TFD are shown by CPK representation with the atoms colored as carbon–violet (FFD) or cyan (TFD), oxygen–red, nitrogen–blue, hydrogen–white (only polar hydrogens of ligand are shown). **h-i**. Docked poses of FFD and TFD in cPLA2, respectively, predicted by IFD. FFD and TFD are shown as ball and stick model with the same atom color scheme. Important amino acids are depicted as sticks with the same color scheme except that carbon atoms are represented in grey. Only polar hydrogens are shown. Dotted yellow lines indicate hydrogen-bonding interactions. Values of the relevant distances are given in Å. **j**. DARTS assay with serial deletion constructs encoding Flag-tagged mutants of cPLA2. cPAL2 (aa 1-750), cPLA2 (aa 126-750), cPLA2 (aa 406-750), cPLA2 (aa 1-479), cPLA2 (aa 1-144). 293T cells were transfected with Flag-tagged mutants of cPLA2 plasmids, as indicated. DARTS assay samples were detected by Flag antibody. **k**. DARTS assay for Ser-505 point mutant of cPLA2. The Ser-505 of cPLA2 was substituted with Ala505. 293T cells were transfected with the point mutant plasmid and DARTS was performed. Point mutated cPLA2 was detected by Flag antibody.

In order to further confirm the associations of cPLA2 with Fexofenadine and Terfenadine, we employed the cellular thermal shift assay (CETSA)^35,36^, which allows for quantification of the change in thermal denaturation temperature of a target protein under different conditions, including those of varying temperature and concentrations of drug of interest. Both Fexofenadine and Terfenadine, particularly Fexofenadine, prevent denaturation of cPLA2 and kept more cPLA2 in the soluble condition under several temperatures, strikingly obvious at 49°C, compared to DMSO (**Fig. 5e, top**). The melt curve showed a significant shift and obvious change of T_m_ in the presence of Fexofenadine and Terfenadine (T_m_ for control DMSO, Terfenadine and Fexofenadine are 46.09, 49.08 and 51.99, respectively) (**Fig. 5e, bottom**). Performance of CETSA at 49°C with different dosages of drugs revealed that Fexofenadine and Terfenadine prevented cPLA2 denaturation in a dose-dependent manner, with the EC50 of 1.025e-007 and 1.449e-009, respectively (**Fig. 5f**).

To further characterize the interactions between Fexofenadine/Terfenadine and cPLA2, we performed both induced-fit docking (IFD) and molecular dynamics (MD) simulations. From IFD simulation, both Fexofenadine and Terfenadine core structures were predicted to stabilize at the MAPK phosphorylation site of cPLA2 at Ser-505 (**Fig. 5g**). Docked poses of Fexofenadine in cPLA2 reveal that the Fexofenadine-binding region was lined by Ser-505 and several residues closed to Ser-505 (**Fig. 5h**). Fexofenadine was predicted to be involved in two hydrogen bonding interactions with Ser-505. The predicted binding-region and hydrogen bonding interaction for Terfenadine were nearly the same as Fexofenadine (**Fig. 5i**). MD simulations, as a complement to IFD simulation, showed that Fexofenadine core structure was majorly stabilized into the binding site predicted by IFD. The protein backbone RMSD deviated up to about 4 Å in the first 3 ns then remained relatively stable till the end of the simulation period, reflecting a relatively stable protein conformation. There was no significant turn-over in the Fexofenadine binding pose as the Fexofenadine RMSD deviated no more than 2 Å since the beginning of simulation (**Fig. S13a**). For the cPLA2-Terfenadine binding complex, both the binding pocket of cPLA2 and the binding pose of Terfenadine showed no significant steric changes with only slight fluctuations on RSMD values after 2 ns (**Fig. S13b**). The monitored 10 ns MD trajectory of cPLA2-Fexofenadine complex are shown in the **Supplemental Video 1**.

### Fexofenadine and Terfenadine inhibit TNF inflammatory activity through binding to the catalytic domain 2 of cPLA2 and the inhibition of the phosphorylation of cPLA2 on Ser-505

cPLA2 contains several domains critical for its functions^37^, including Ca^2+^ binding domain (C2D), catalytic domain 1 (CD1) and catalytic domain 2 (CD2) (**Fig. 5j**). We sought to identify the domain by which Fexofenadine/Terfenadine targets to cPLA2. For this purpose, we generated serial N-terminal and C-terminal deletion mutants and tested their interactions with Fexofenadine by use of DARTS (**Fig. 5j**). Similar to the protective effect seen with intact cPLA2, Fexofenadine retained protective effects for mutants with N-terminal C2D deletion (i.e. cPLA2 (126-750) and further deletion of CD1 (i.e. cPLA2(406-750)), indicating that CD2 domain is the binding domain of Fexofenadine. Indeed, Fexofenadine did not show any protective effects to the mutants lacking CD2 domain (i.e. cPLA2 (1-479) and cPLA2 (1-144)). Collectively, these sets of assays identify the CD2 domain of cPLA2 as the binding domain of Fexofenadine and Terfenadine that is metabolized into Fexofenadine *in vivo*.

Interestingly, both IFD and MD simulations indicated that Ser-505, which is located in the CD2 domain of cPLA2, is the critical amino acid for the interactions between Fexofenadine/Terfenadine and cPLA2. We next determined whether the substitution of Ser-505 with Ala through the site-directed mutagenesis affected the binding of Fexofenadine to cPLA2. DARTS assay clearly demonstrated that Fexofenadine lost its protective effect on this point mutant of cPLA2 (**Fig. 5k**), further demonstrating that Ser-505 is the critical amino acid required for Fexofenadine targeting to cPLA2.

It is well established that the phosphorylation of cPLA2 on Ser-505 by upstream kinases p-p38 and p-REK1/2 is required for its enzymatic activity^38^. Since Ser-505 is an essential amino acid of the binding site for Fexofenadine targeting to cPLA2, we next examined whether Fexofenadine affected TNF-activated phosphorylation of cPLA2 on Ser-505 (**Fig. 6a**). As expected, TNF-α activated the phosphorylation of p38 and ERK1/2 as well as cPLA2 in BMDMs. Fexofenadine treatment did not affect the phosphorylation of p38 or ERK1/2, but abolished the phosphorylation of cPLA2 on Ser-505 (**Fig. 6a**), further indicating that Fexofenadine directly targets to cPLA2, without affecting its upstream mediators in the TNF-activated cPLA2 inflammatory pathway.

**Fig. 6.**
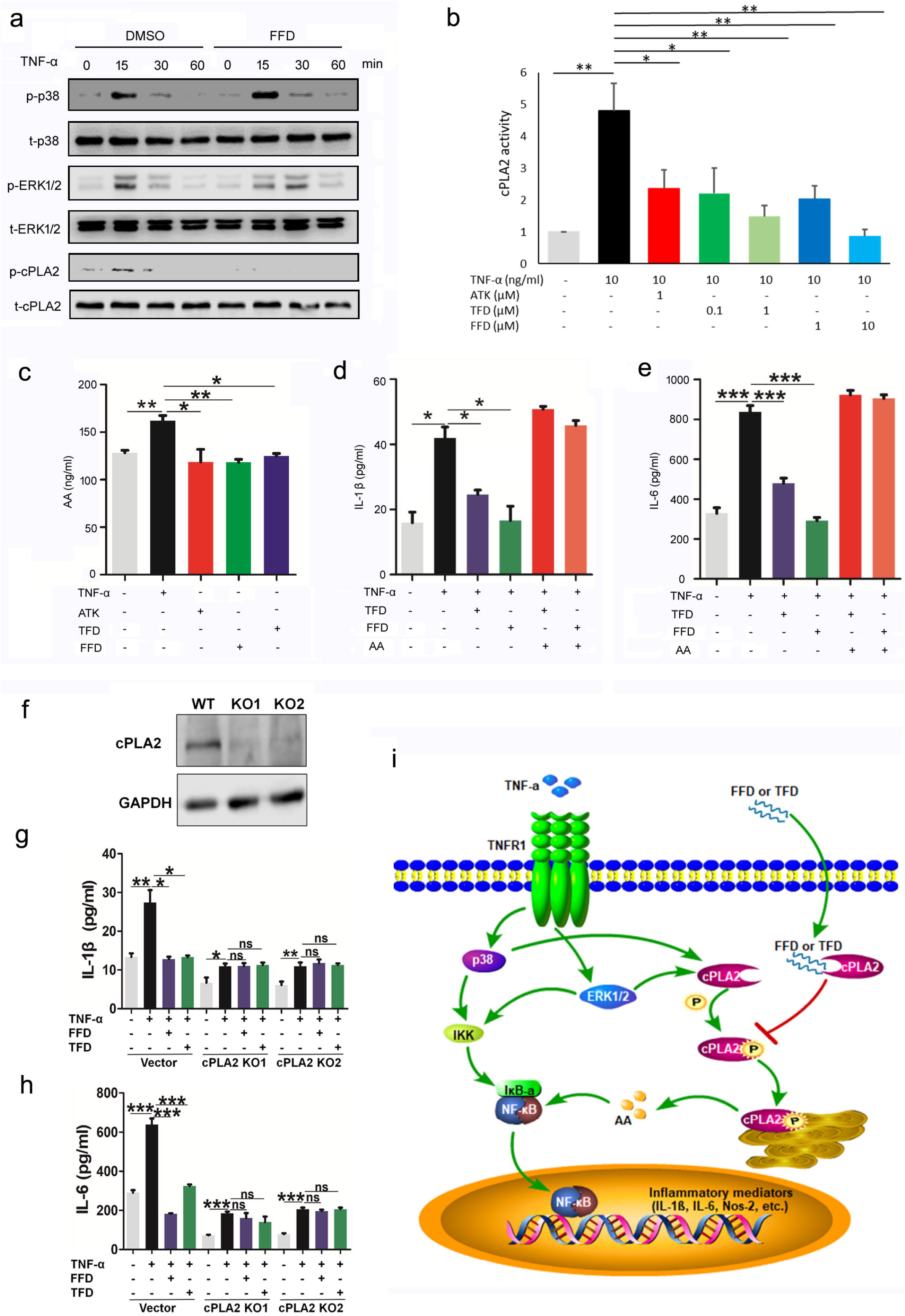
Fexofenadine and Terfenadine inhibit TNF activity through binding to the catalytic domain 2 of cPLA2 and inhibition of the phosphorylation of cPLA2 on Ser-505. **a**. Fexofenadine (FFD) inhibits the phosphorylation of cPLA2 on Ser-505. BMDM cells were treated with TNF-α (10 ng/ml) in the absence or presence of FFD (10 μM) for various time points, as indicated. p-p38, t-p38, p-ERK1/2, t-ERK1/2, p-cPLA2 (specifically for phosphorylated Ser-505), t-cPLA2 were detected by Western blot with corresponding antibodies. **b**. Fexofenadine/Terfenadine (TFD) inhibits TNF-induced cPLA2 activity in living cells. RAW264.7 cells transfected with an expression plasmid encoding cPLA2 were treated with TNF-α and ATK, or TFD, or FFD overnight. Cells lysate was used for cPLA2 activity analysis. **c**. FFD/TFD inhibits TNF-induced arachidonic acid (AA) production. The AA levels in BMDMs without or with TNF-α (10 ng/ml) in absence or presence of FFD or TFD for 48 hours were examined using a commercial ELISA kit. ATK was used as a positive control. **d-e**. Addition of AA abolished FFD/TFD inhibition of TNF-induced cytokine release. BMDMs were treated with TNF-α (10 ng/ml), AA (10 μM), and FFD (10 μM)/TFD (1 μM), as indicated. The levels of IL-1β and IL-6 were detected by ELISA. **f**. Knock out efficiency of cPLA2 using CRISPR-Cas9 technique in RAW264.7 cells, assayed by Western blot. Two individual knockout clones (KO1 and KO2) were employed. **g-h**. Deletion of cPLA2 abolished FFD/TFD inhibition of TNF-induced cytokine release. WT and cPLA2 KO RAW264.7 cells were treated without or with TNF-α (10 ng/ml) in absence or presence of FFD (10 μM)/TFD (1 μM) for 48 hours. The levels of IL-1β and IL-6 in medium were detected by ELISA. (* p<0.05, ** p<0.01, ***p<0.001). i. A proposed model for explaining the anti-TNF activity of FFD/TFD through targeting cPLA2 pathway.

As mentioned earlier, the phosphorylation of cPLA2 on Ser-505 is required for its enzymatic activity, we next examined whether Fexofenadine/Terfenadine affected the activity of cPLA2. Similar to arachidonyl trifluoromethyl ketone 27 (ATK), a known cPLA2 inhibitor and serving as a positive control, both Fexofenadine and Terfenadine completely abolished TNF-α induced cPLA2 activity and their inhibitions on cPLA2 activity are dosage-dependant (**Fig. 6b**).

Inflammatory conditions, including elevated TNF-α, promote the cPLA2 translocation to intracellular phospholipid membrane. The major function of cPLA2 is to promote phospholipid hydrolysis to produce arachidonic acid (AA)^39^, which, in turn, actives NF-κB-mediated inflammation^23^. Consequently, we ascertained whether Fexofenadine and Terfenadine inhibited TNF-α induced AA production, the data in **Fig. 6c** revealed that this was the case. Then we examined whether Fexofenadine and Terfenadine inhibition of AA production contributed to their inhibition of TNF-α induced inflammation. As shown in **Fig. 6d-e**, supplementation of medium with AA completely eliminated the inhibitory influence of Fexofenadine/Terfenadine upon TNF-α induced release of inflammatory cytokines IL-1β and IL6, suggesting that inhibition of AA production by Fexofenadine and Terfenadine is important for their anti-TNF-α activity.

We next sought to further determine the importance of cPLA2 in mediating Fexofenadine and Terfenadine’s anti-TNF-α activity. We deleted this gene using CRISPR-Cas9 technique (**Fig. 6f**). This technology produced near complete deletion of cPLA2 (**Fig. 6f**). Importantly, Fexofenadine and Terfenadine mediated inhibition of TNF-α activity was entirely or almost entirely lost in cPLA2 knockout cells, respectively (**Fig. 6g-h**). Taken together, these findings indicated that cPLA2 is required for Fexofenadine and Terfenadine mediated anti-TNF-α activity.

## DISCUSSION

TNF-α signaling associates with various pathophysiological processes and enormous efforts have been devoted to develop treatments for TNF-α associated diseases and conditions^7,9^. In this study we performed three rounds of screenings of an FDA-approved drug library, and isolated Fexofenadine and Terfenadine, two selective histamine receptor 1 antagonists, as two TNF-α inhibitors (**Figs. S1-3**). Comprehensive evidences, including RNA-seq, transcription factor enrichment analysis, downstream cytokine expression and release, NF-κB nuclear translocation and activity, osteoclastogenesis as well as *in vivo* reporter and transgenic mice, validated their anti-TNF-α activities (**Figs. 1, S4-6**). Subsequent *in vivo* animal models, including TNF-tg, collagen induced arthritis, DSS- and TNBS-induced colitis demonstrated that Fexofenadine and Terfenadine are therapeutic against these autoimmune disorders, better than, or at least as good as, the current small molecule drugs for treating rheumatoid arthritis and inflammatory bowel diseases (**Figs. 2–4**). Although both Fexofenadine and Terfenadine yield comparable effects in inhibiting TNF-α activity *in vitro* and in the disease models tested, we consider Fexofenadine to be the more promising drug for treating various TNF-α associated conditions, because Terfenadine has been suspended due to the potential adverse events^40^, whereas Fexofenadine, the major active metabolite of Terfenadine, has proven to be safer and does not have significant risks associated with Terfenadine treatment^41^. Due to its safety and efficacy, Fexofenadine is widely used as a non-prescription medicine, sometimes called an over-the-counter (OTC) drug, for treating various allergic diseases.

Currently marketed TNF-α blockers, such as etanercept (Enbrel) and adalimumab (Humira), have a demonstrated record of safety and efficacy in the treatment of autoimmune diseases. However, Fexofenadine demonstrates features that suggest it may compare favorably to these established biologics. For example, all currently marketed anti-TNF therapies bind to TNF-α and inhibit its binding to TNF receptors; in contrast, Fexofenadine targets the downstream cPLA2 mediator, but not the upstream, of TNF-α signaling. In addition, it also targets H1R1. Due to this alternate mechanism of action, Fexofenadine may be effective for up to ~50% of patients who fail to respond to current TNF-α blockers^12^. As a well-tolerated and generically available oral OTC drug, Fexomenadine’s safety, convenience, and cost-effectiveness suggest that it may be an attractive and viable agent for the clinical treatment of inflammatory autoimmune diseases, particularly rheumatoid arthritis and inflammatory bowel diseases in which Fexofenadine has been proven to be effective in the preclinical animal models (**Figs. 2–4, S7-9**).

Fexofenadine is known to be a highly selective antagonist to H1 receptor 1^30,42^. Surprisingly, suppression of H1 receptor 1 does not affect Fexofenadine-mediated anti-TNF-α activity (**Fig. S10**). In addition, additional 7 known H1R1 inhibitors do not have anti-TNF activity (**Fig. S11**). Although Fexofenadine potently inhibits TNF-α signaling *in vitro* and *in vivo*, it does not affect the binding of TNF-α to its receptors and cell surface, clearly different from clinically used TNF-α inhibitors (**Fig. S12**). Excitingly, through combined use of drug affinity responsive target stability assay, proteomics, cellular thermal shift assay, induced-fit docking and molecular dynamics, we identified cPLA2 as a previously unrecognized target of Fexofenadine. Fexofenadine binds to catalytic domain 2 and inhibits the phosphorylation of cPLA2 on Ser-505. Further, deletion of cPLA2 abolished Fexofenadine inhibition of TNF induced AA production and downstream cytokine release (**Figs. 5–6**). A proposed model for explaining the anti-TNF activity of Fexofenadine through directly targeting cPLA2 pathway is shown in **Fig. 6i**. TNF-α binds to TNFR1 and activates p38 and ERK1/2, followed by the phosphorylation of cPLA2 on Ser-505. Phosphorylated cPLA2 then translocates from the cytosol to hydrolyze membrane phospholipids, leading to the production of AA. AA, in turn, actives NF-κB, leading to cytokine release and inflammation. Dissimilar to current TNF inhibitors that disturb TNF/TNFR interactions at the initiation of the signaling cascade, Fexofenadine diffuses into the cells and directly binds to cPLA2 and inhibits its phosphorylation on Ser-505, followed by the inhibition of cPLA2/AA/ NF-κB inflammatory pathway.

TNF-α regulation of immune cells, such as macrophage polarization and differentiation of T cell populations, is a known component in the pathogenesis of autoimmune diseases^7^. We found that both Fexofenadine and Terfenadine dose-dependently inhibited the expressions of inflammatory M1 macrophage-associated genes *Il6* and *Nos2* (**Fig. S14a**), while markedly increased the expressions of anti-inflammatory M2 macrophage-associated genes *Arg1* and *Mgl1* (**Fig. S14b**). Furthermore, Fexofenadine and Terfenadine significantly suppressed the differentiation of IFN □-positive Th1 subpopulation. Fexofenadine also significantly inhibited Th17 cell differentiation, whereas negligible effects were observed with regard to the differentiation of Th2 and regulatory T cells (Tregs) (**Fig. S15a-d**). It is expected that Fexofenadine regulation of macrophages and T cells may also contribute to its anti-TNF/cPLA2 activity *in vivo*, which warrant further investigations. Additionally, although Fexofenadine is therapeutic against inflammatory arthritis spontaneously developed in TNF-tg mice (**Fig. 2**), and Fexofenadine-mediated anti-TNF-α activity depends on cPLA2 (**Fig. 6f-h**), but not histamine H1 receptor (**Fig. S10**), its anti-histaminic action may also contribute to its therapeutic effects in autoimmune diseases.

Similar to TNF-α, cPLA2 is also known to play an important role in regulating autoimmune diseases^43,44^. cPLA2 is implicated in synovitis and joint destruction in rheumatoid arthritis by regulating the production of inflammatory mediators^45^. cPLA2 up-regulation leads to a greater disease severity phenotype in the DSS-induced colitis mouse model, whereas intravenous administration of antisense oligonucleotides against cPLA2 prevents disease development, suppresses NF-κB activation, and decreases expressions of pro-inflammatory proteins^46^, indicating the potential treatment target role of cPLA2 in colitis. Interestingly, Fexofenadine was reported to have potential treatment effects in the murine DSS-induced inflammatory bowel disease model through an unclear mechanism^47^. Our data demonstrating that Fexofenadine directly targets to cPLA2/AA and is therapeutic against inflammatory bowel disease link these previous reports together nicely and provide a mechanistic foundation for previous findings and future work.

In addition to autoimmune diseases, cPLA2 is also involved in the pathogenesis of many other disorders, particularly neurodegenerative diseases, cardiovascular diseases and cancer^48–55^. Intense efforts have been invested to identify potent cPLA2α inhibitors in the past several decades, unfortunately, toxicity and the poor absorption of anti-cPLA2 compounds from intestine were the significant challenges for clinical application, although numerous cPLA2 inhibitors have been tested in clinical trials^56,57^. Our findings that Fexofenadine, an OTC drug widely used for treating allergic diseases, targets and inhibits cPLA2 and is effective in various animal models of autoimmune diseases may provide innovative interventions to overcome the current bottlenecks in the efforts to develop cPLA2 targeting treatments.

In summary, this study identifies Fexofenadine as an inhibitor of TNFα signaling, and uncovers a new strategy for inhibiting this cardinal pathway of inflammation. In addition, this study also identifies cPLA2 as a new target of Fexofenadine, thus providing new insights into the understanding of Fexofenadine’s action and underlying mechanisms, and a solid foundation for future discoveries relating to this Fexofenadine/cPLA2 interaction. Further, it also identifies Fexofenadine as a novel antagonist of cPLA2, suggesting that Fexofenadine can also be used for treating various cPLA2 associated diseases, including autoimmune diseases, neurodegenerative diseases, cardiovascular diseases and cancer. With the consideration that both TNF-α and cPLA2 are involved in a plethora of disease processes, the identification of Fexofenadine as an inhibitor of both TNF-α and cPLA2, and manipulation of new antagonist of the TNF/cPLA2 pathway may lead to innovative therapeutics for various pathologies and conditions, significantly broadening the application of this OTC drug in addition to allergic diseases.

## METHODS SUMMARY

### *In vitro* and *in vivo* screen of FDA-approved drug library

NF-κB-*bla* THP-1 cell line in which NF-κB beta-lactamase reporter gene was stably integrated, RAW 264.7 macrophages in which NF-κB luciferase reporter gene was transiently transfected, and TNF-tg: NF-κB-Luc double mutant reporter mice were employed to screen the drug library.

### *In vitro* and *in vivo* assays for examining the blockade of TNF actions by Fexofenadine and Terfenadine

RNA-seq, transcription factor enrichment analysis, downstream cytokine expression and release, NF-κB translocation and activity, osteoclastogenesis with bone marrow derived macrophages and RAW264.7 macrophages, as well as in vivo reporter mice.

### *In vivo* assays for defining the anti-inflammatory activity of Fexofenadine and Terfenadine using various animal models

Administration of Fexofenadine, Terfenadine or clinically used positive controls into TNFα transgenic (Tg) mice, collagen-induced arthritis of DBA/1 mice, DSS- and TNBS-induced inflammatory bowel diseases of C57/BL6 mice.

### Identification and characterization of the binding of Fexofenadine/Terfenadine to cPLA2

Drug affinity responsive target stability assay, proteomics, cellular thermal shift assay, information field dynamics and molecular dynamics; Solid-phase binding and flow cytometry was used to examine the effects of Fexofenadine/Terfenadine on TNF/TNFR interactions.

### Assays for examining the inhibition of cPLA2 phosphorylation and activity by Fexofenadine/Terfenadine as well as the dependence on cPLA2

Phosphorylation of p38, Erk1/2 and cPLA2 by TNF-α and their inhibition by Fexofenadine/Terfenadine, activation of cPLA2 activity and AA production by TNF-α and their inhibition by Fexofenadine/Terfenadine, dependence on the presence and activity of cPLA2 and AA production of Fexofenadine/Terfenadine’s inhibition of TNF-α, were determined.

<u>Detailed experimental procedures are provided in Supplementary Materials</u>

## Acknowledgements

This work was supported partly by NIH research grants R01NS103931, R01AR062207, R01AR061484, and a DOD research grant W81XWH-16-1-0482.

Patents have been filed by NYU School of Medicine that claim Fexofenadine and Terfenadine Target Cytosolic Phospholipase A2. U.S. Application No. 62/701,806

