## Supplementary Data for "Fexofenadine inhibits TNF signaling through targeting to cytosolic phospholipase A2 and is therapeutic against autoimmune diseases"

This Supplementary Materials file includes:

**Figs. S1 to S15** (Pages 2-16)

**Experimental Procedures** (Pages 17-31)

**References** (Pages 32-33)

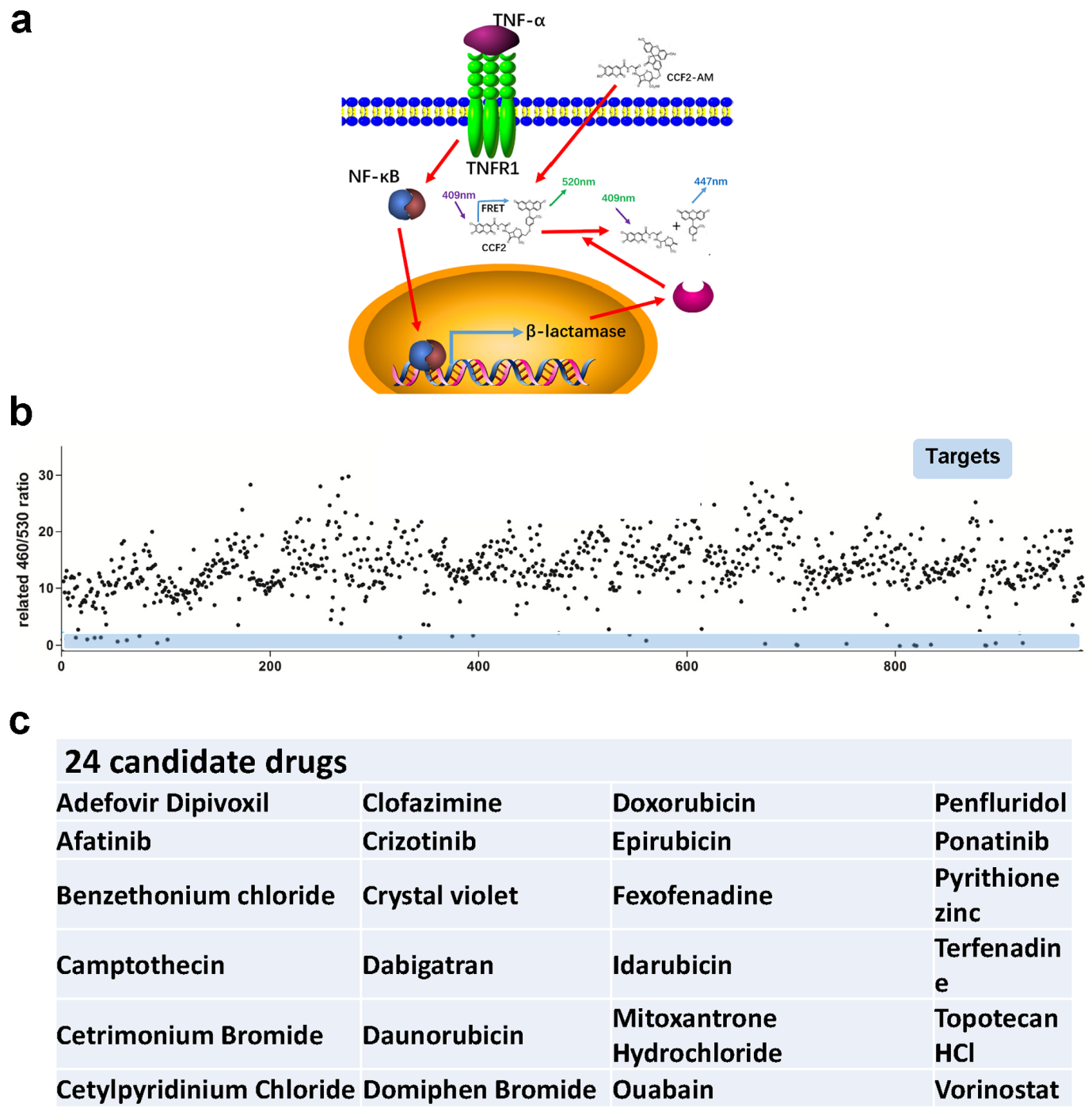

**Fig. S1. The 1st round screen of FDA approved drugs.** THP-1 cell line with NF-κB beta-lactamase reporter gene stably integrated was used. The first screening was repeated 3 times. **a.** Schematic of the first screening using THP-1 cell line. Cells were incubated overnight with drugs (10 μM) followed by incubation with TNF-α (10ng/ml) for 6 hours. Beta-lactamase reporter gene activity was measured. **b.** The analysis of the first round screen. **c.** Identities of twenty-four candidate drugs isolated from 1st round screen.

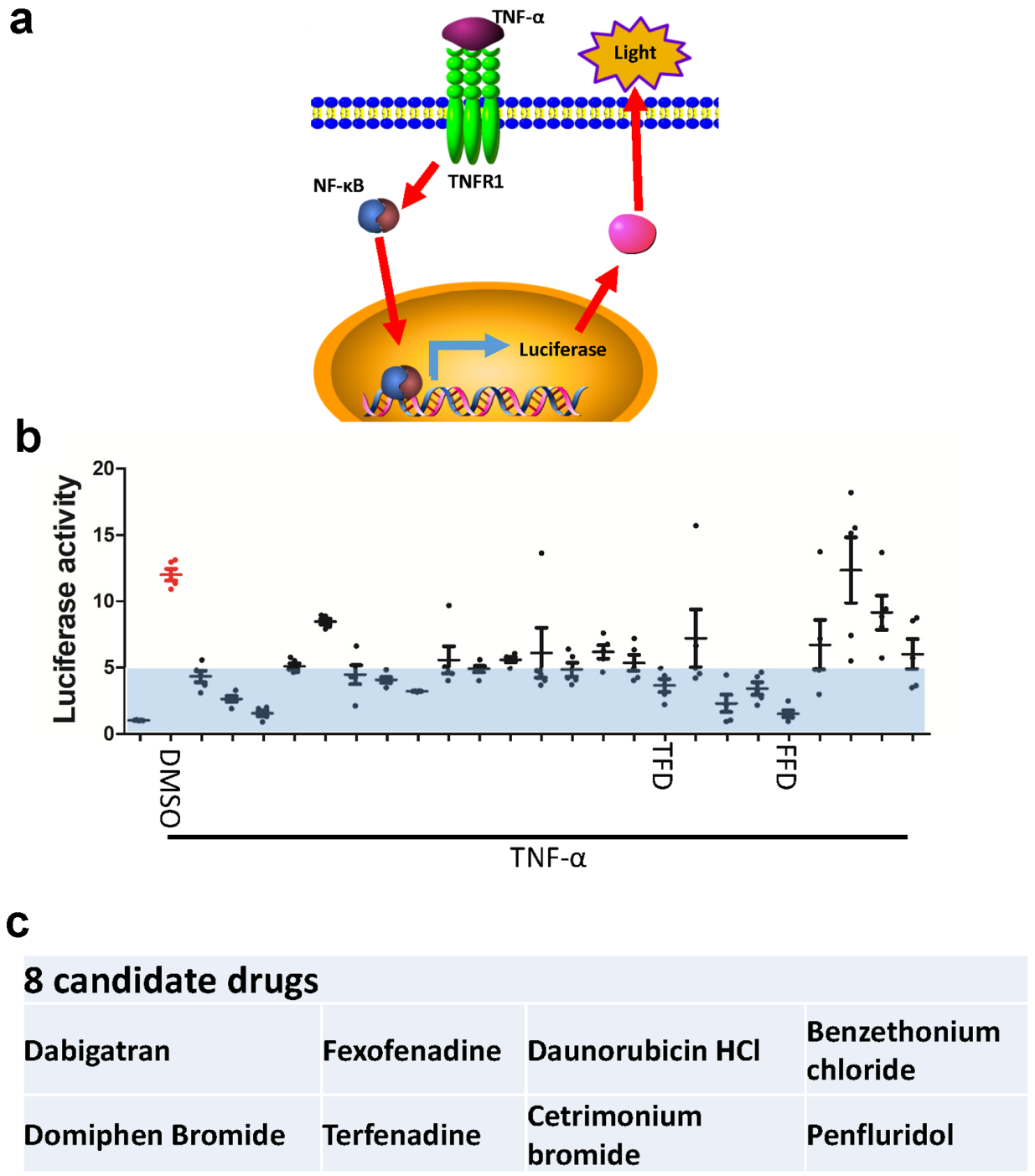

**Fig. S2. The 2nd round screen of FDA approved drugs. a.** The diagram of the second screening. NF-κB luciferase reporter plasmid was transfected into RAW 264.7 cells to confirm the results of the first round screening. Cells were treated with drugs (10μM) overnight, followed by treatment with TNF-α (10ng/ml) for 6 hours, bioluminescence was taken as a measure of luciferase activity. **b.** The analysis of the second screening. TFD and FFD indicate Terfenadine and Fexofenadine, respectively. **c.** Identities of eight candidate drugs isolated from 2nd round screen.

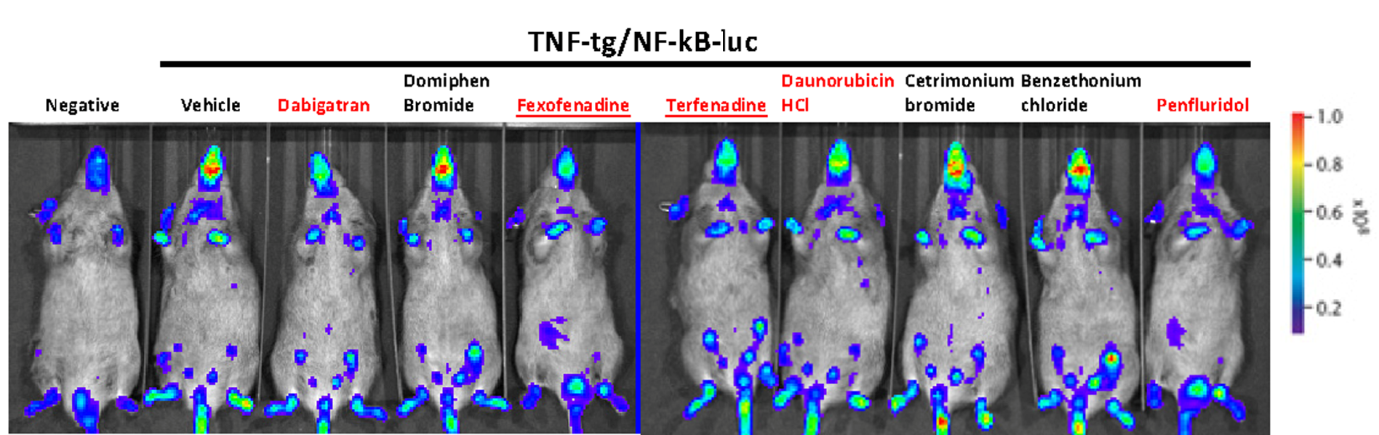
**Fig. S3. The 3rd round screen of FDA approved drugs.** TNF-tg:NF-κB-Luc mice were generated to confirm the *in vivo* activity of 8 drugs isolated in preliminary screens. After treatment for 7 days with indicated compounds, the luciferase reporter signal was detected by IVIS. The drugs that show anti-TNF activity *in vivo* are highlighted in Red. Terfenadine and Fexofenadine are underlined.

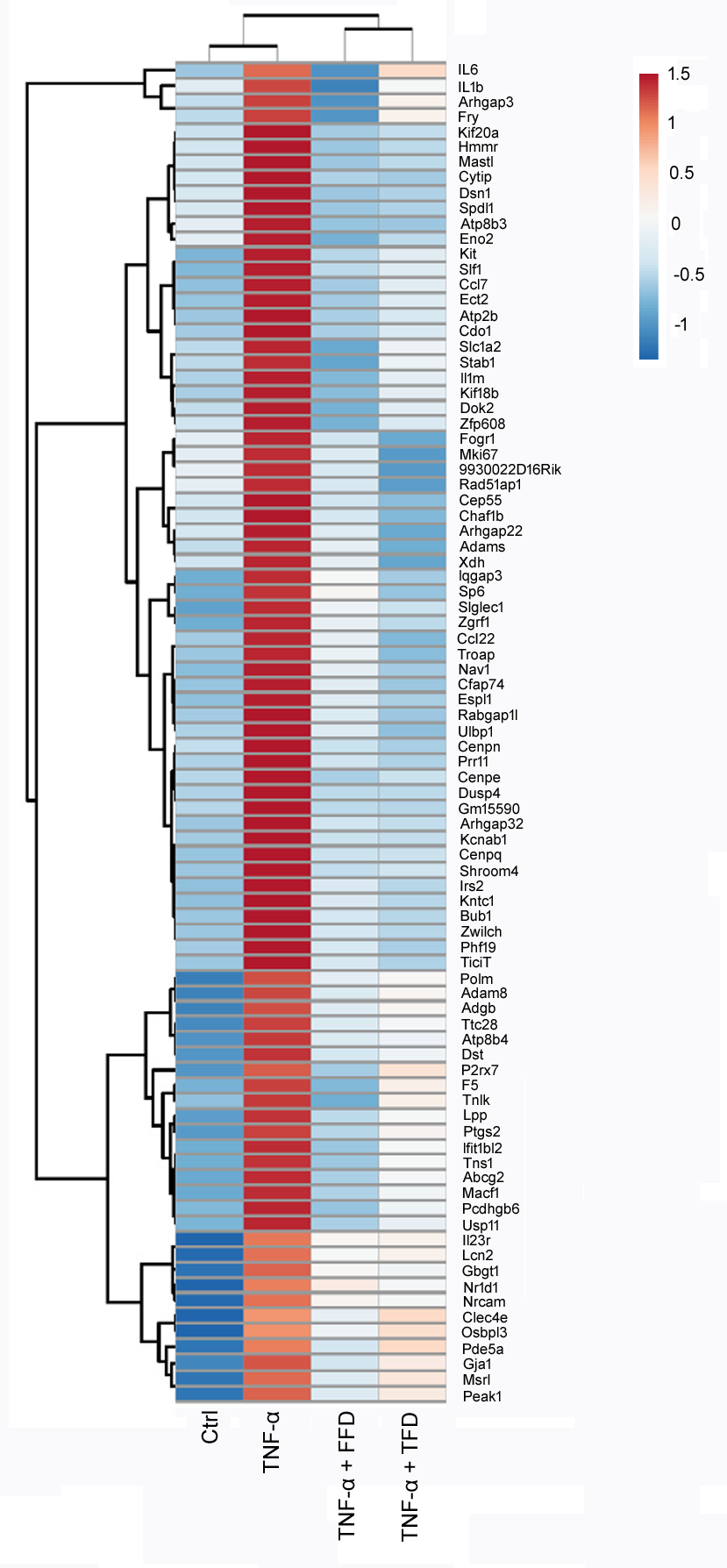

**Fig. S4. RNAseq analysis.** BMDMs were treated without (Ctrl) or with TNF-α (10ng/ml) in absence or presence of Fexofenadine (FFD) (10 μM) or Terfenadine (TFD) (1 μM) for 24 hours. Total RNA were extracted for RNAseq. The genes which were induced by TNF-α and inhibited by both Terfenadine and Fexofenadine were summarized and analyzed.

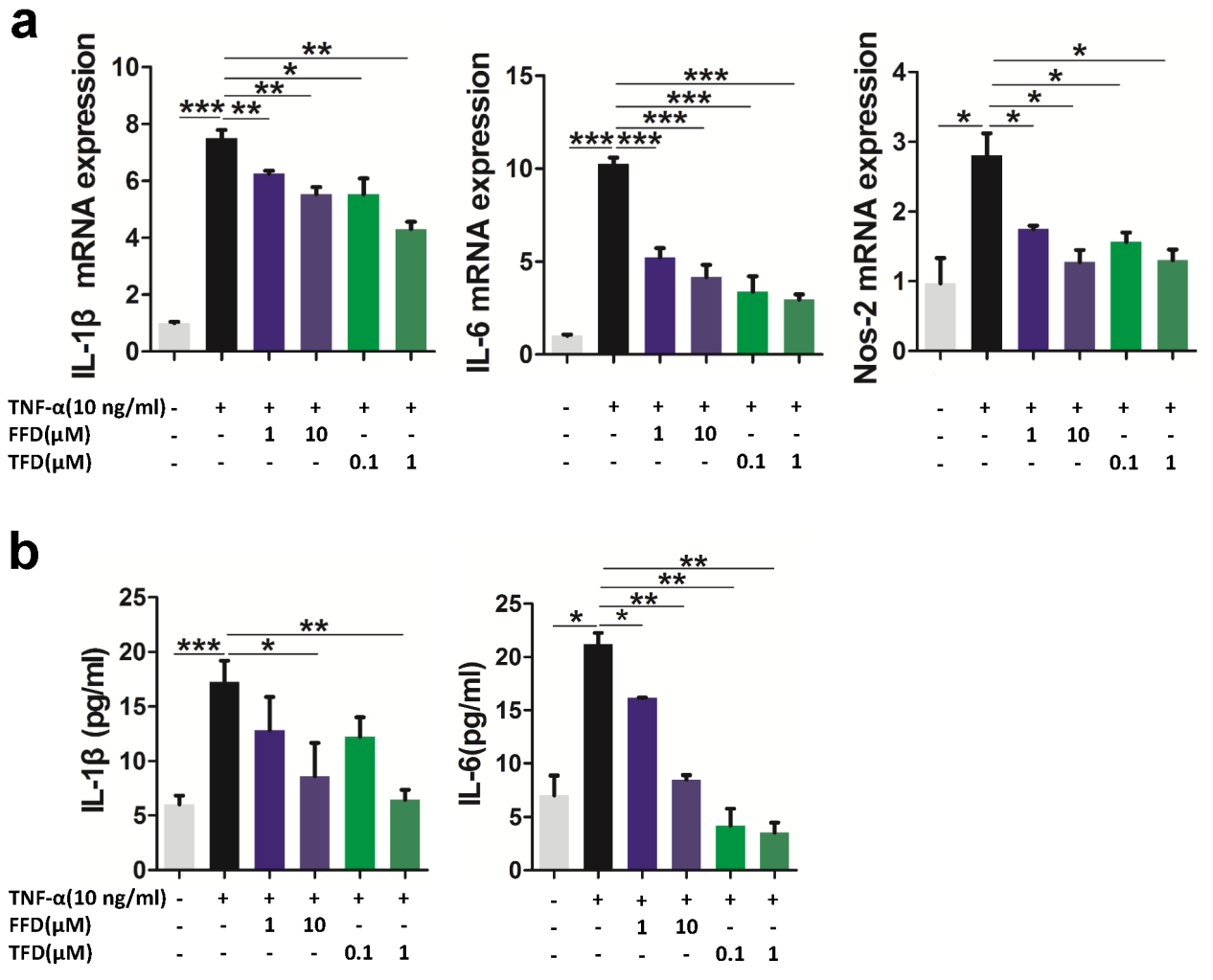

**Fig. S5. Terfenadine and Fexofenadine inhibit TNF-α activity in RAW264.7 cells.** RAW264.7 cells were incubated with TNF-α(10 ng/ml) in the absence or presence of various dosages of Terfenadine (TFD) or Fexofenadine (FFD), as indicated, for 24 hours prior to collection for real time PCR or for 48 hours for ELISA. **a.** The mRNA level of IL-1β, IL-6 and Nos-2 were detected by qRT-PCR. **b.** The secretion level of IL-1β and IL-6 was tested by ELISA. (* p<0.05, ** p<0.01, ***p<0.001)

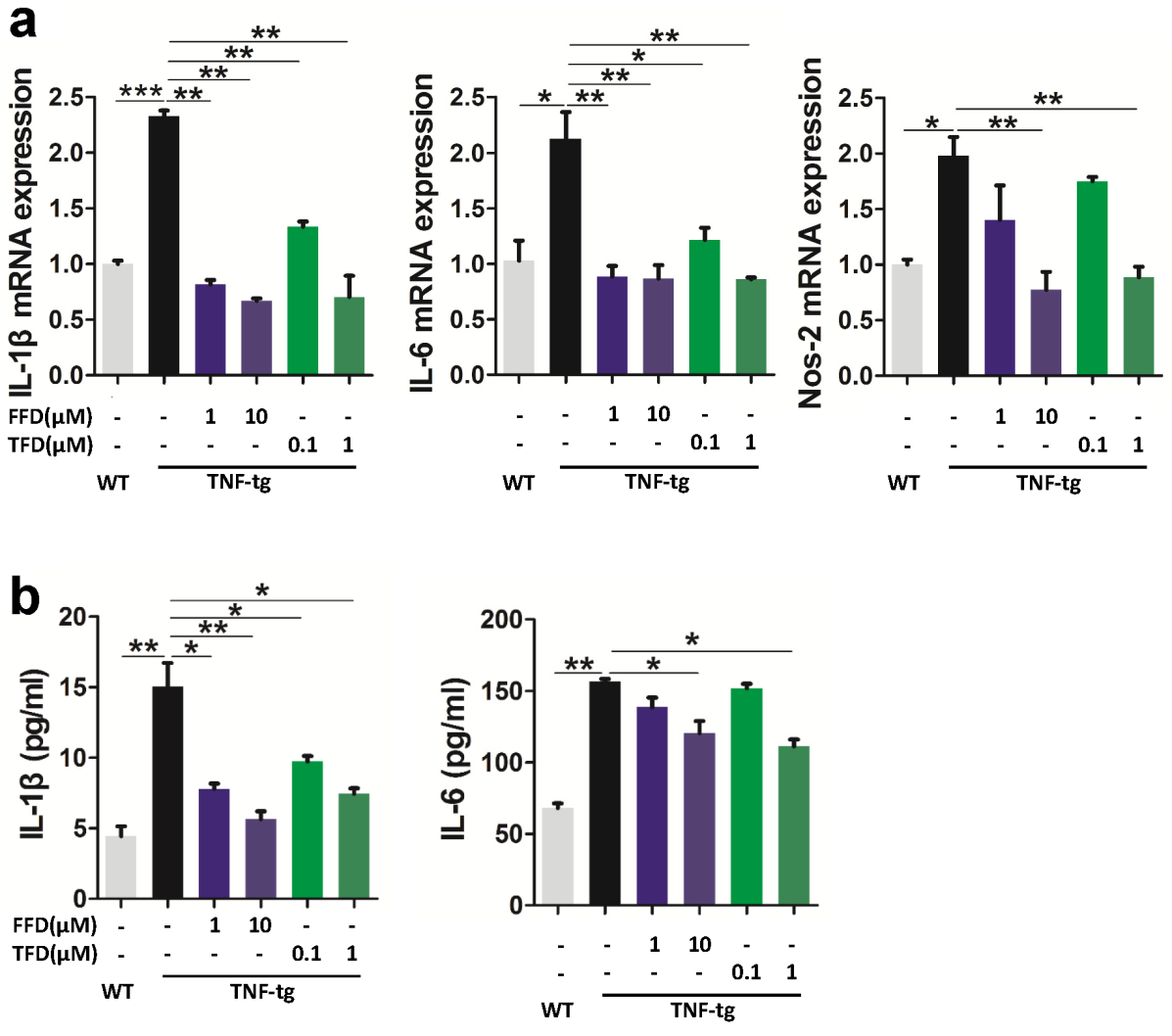

**Fig. S6. Terfenadine and Fexofenadine exhibit anti-TNF activity in BMDM cells isolated from TNF transgenic mice.** BMDMs were first incubated with M-CSF (10 ng/ml) for 6 days, Fexofenadine (FFD, 1μM, 10μM) or Terfenadine (TFD, 0.1μM, 1μM) were added for 24 hours for real time PCR or 48 hours for ELISA. **a.** The mRNA expression of IL-1β, IL-6 and Nos-2 were detected by qRT-PCR. **b.** The levels of IL-1β and IL-6 in medium were tested by ELISA. (* p<0.05, ** p<0.01, ***p<0.001)

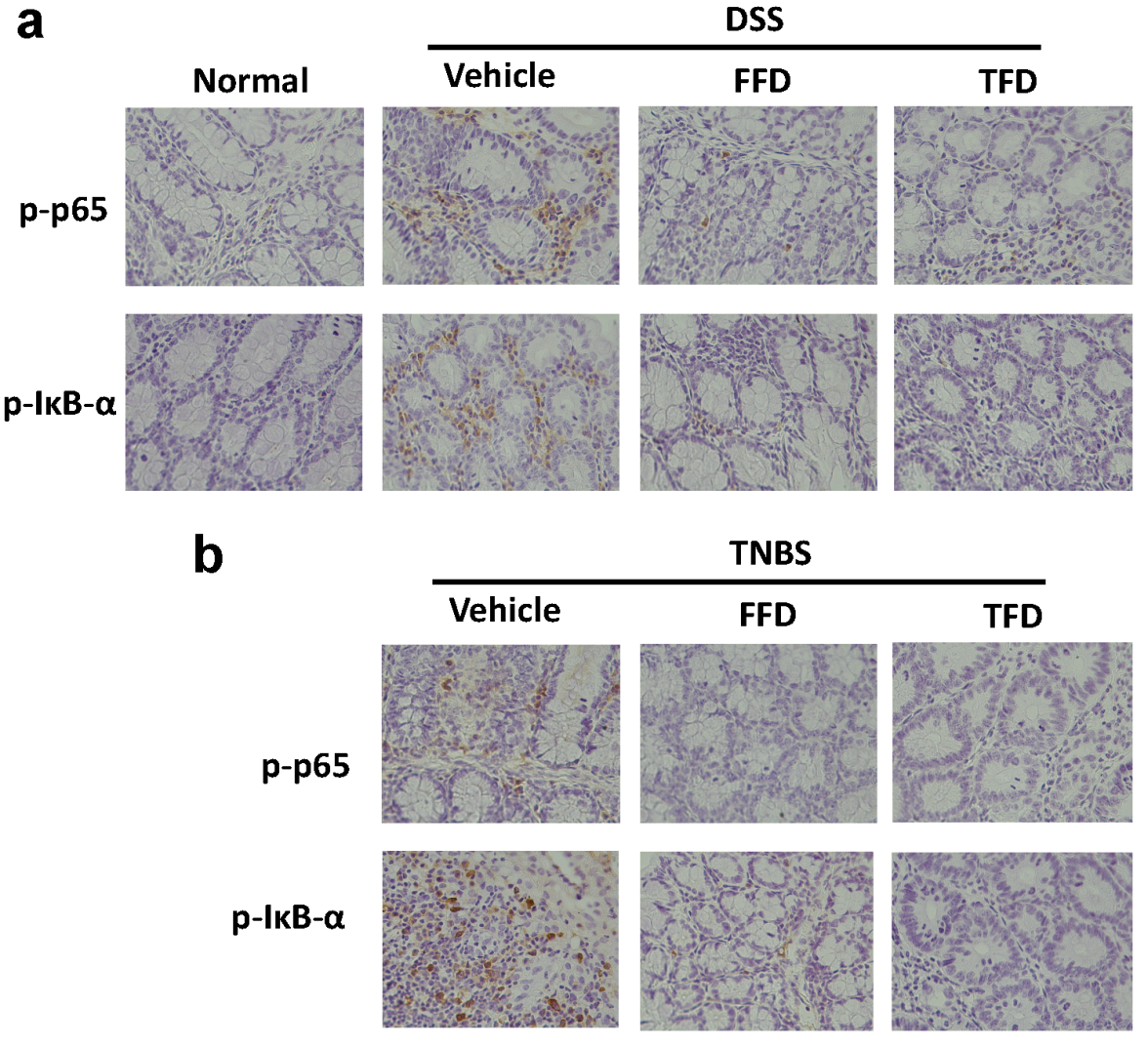

**Fig. S7. Immunohistochemistry (IHC) analysis of p-p65 and p-IκB-α in colon tissues. a.** IHC staining of p-p65 and p-IκB-α in paraffin embedded samples collected from DSS model mice. **b.** IHC staining of p-p65 and p-IκB-α in paraffin embedded samples collected from TNBS model mice. Terfenadine and Fexofenadine are indicated with TFD and FFD, respectively.

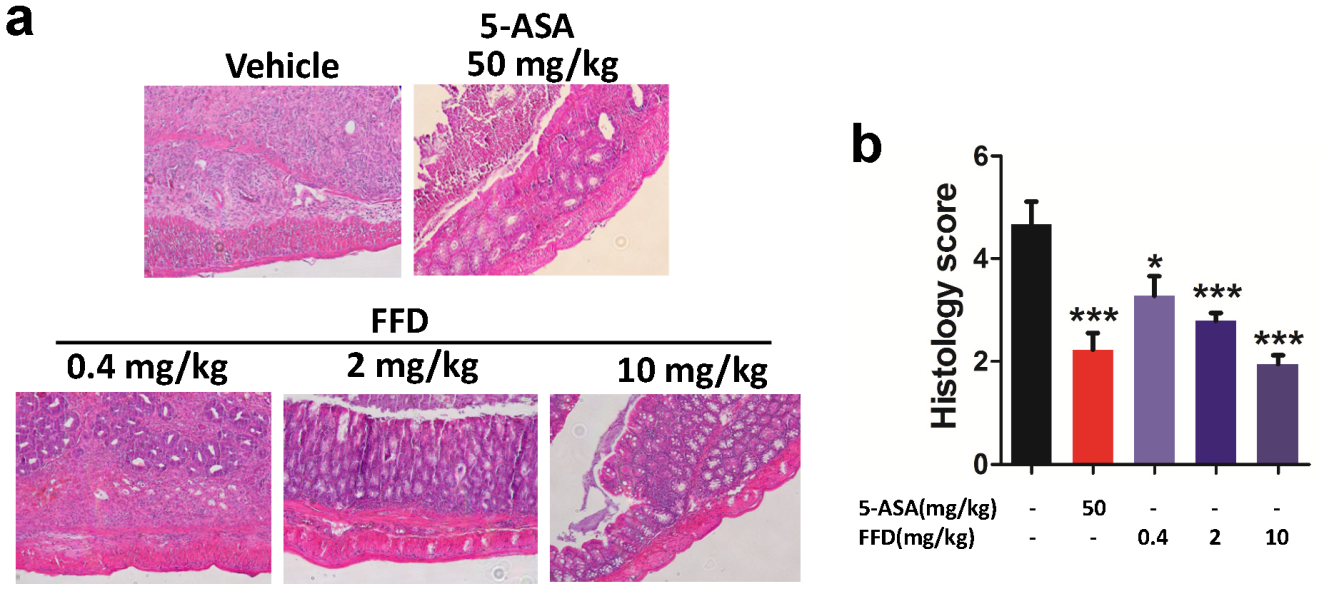

**Fig. S8. Histological staining and histological score of Fexofenadine** (**FFD)-treated and controls TNBS model mice. a.** H&E staining was performed to visualize inflammation of colon in the TNBS model. **b.** The quantification of histological score. (* p<0.05, ***p<0.001)

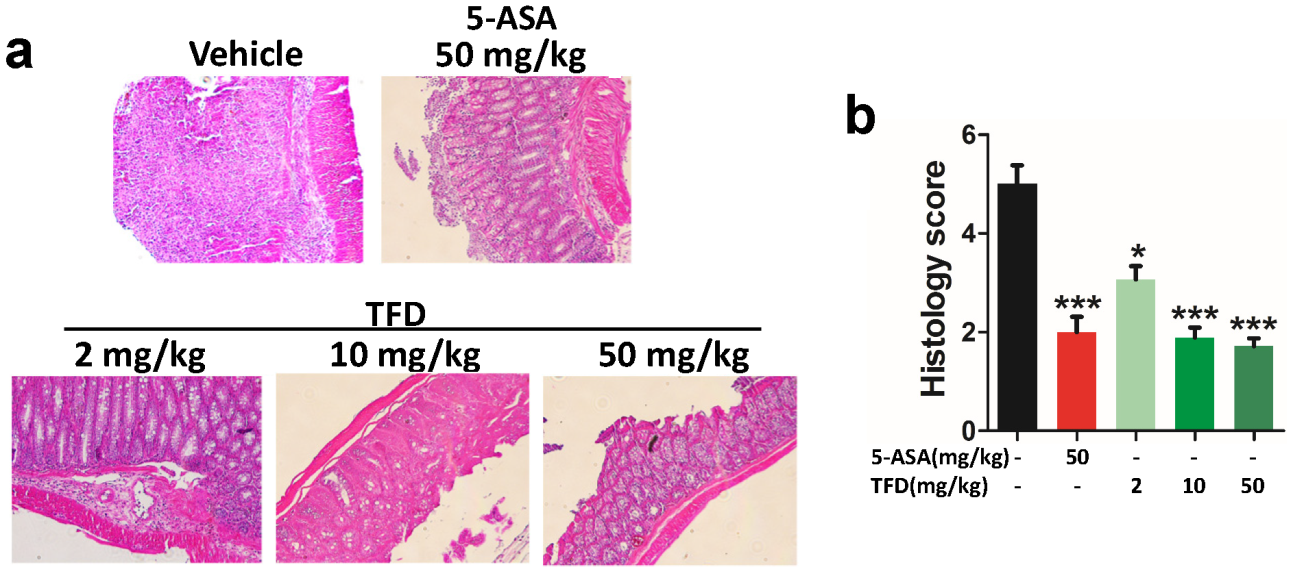

**Fig. S9. Histological staining and score of Terfenadine (TFD)-treated and controls TNBS model mice. a.** The inflammation of colon in TNBS model was detected by H&E staining. **b.** The quantification of histological score. (* p<0.05, ***p<0.001)

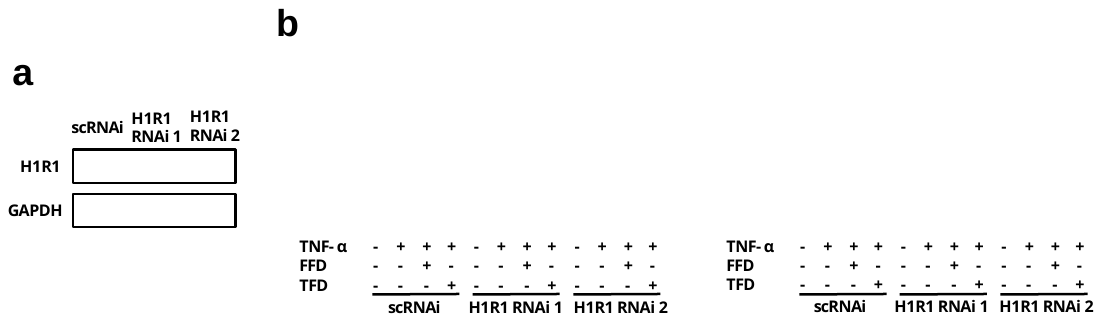

**Fig. S10. The anti-TNF activity of Terfenadine (TFD) and Fexofenadine (FFD) does not depend on H1R1. a.** Immunoblotting analysis to examine the knockdown efficacy of siRNA against H1R1**. b.** RAW264.7 cells transfected with scrambled control siRNA (scRNAi) or H1R1 RNAi were treated with or without TNFα (10ng/ml) in absence or presence of FFD (10μM)/TFD(1μM) for 48 hours**.** The levels of IL-1β and IL-6 in the medium were detected by ELISA (* p<0.05, ** p<0.01, ***p<0.001)

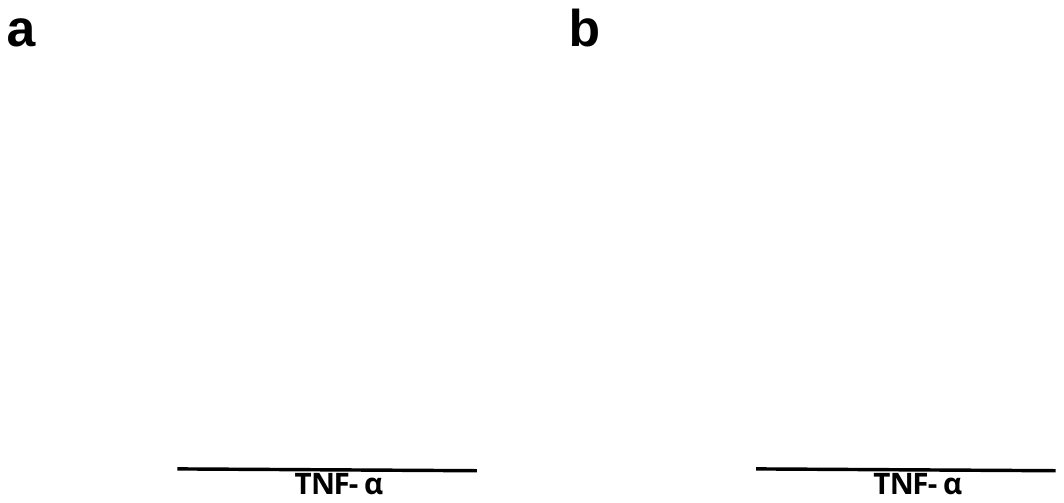

**Fig. S11. Comparison of The anti-TNF activity between Terfenadine (TFD) and Fexofenadine (FFD)and other known H1R1 inhibitors.** BMDM cells were treated without or with TNF-α (10ng/ml) in absence or presence of various H1R1 inhibitor, as indicated, for 48 hours. The levels of IL-1β and IL-6 in medium were detected by ELISA.

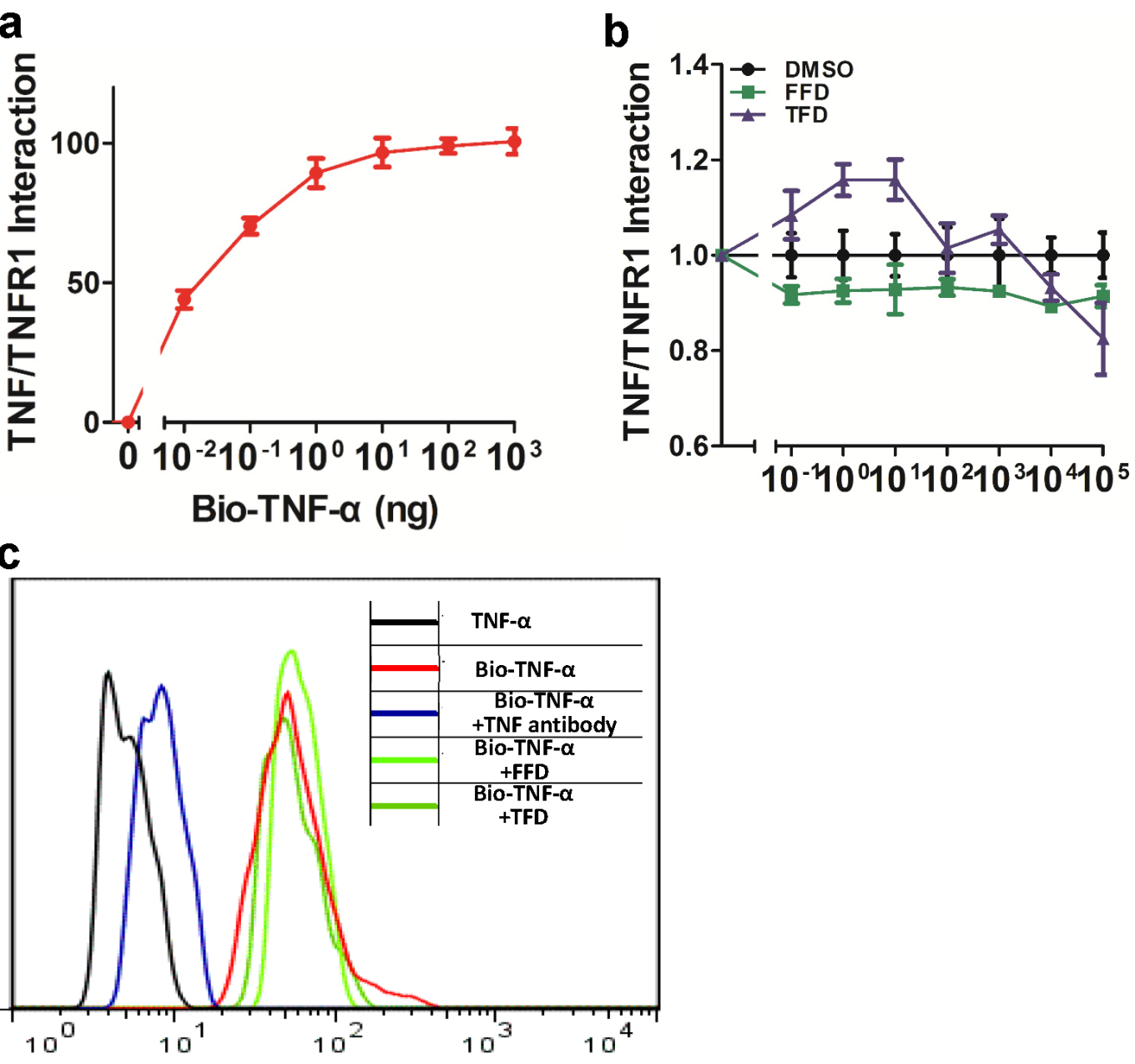

**Fig. S12. Terfenadine (TFD) and Fexofenadine (FFD) do not affect the binding of TNF-α and TNFR1 and to the cell surface. a.** Solid phase binding was used to reveal the dose-dependent binding of TNF-α to TNFR1. **b.** The binding of TNF-α to TNFR1 in the presence of DMSO (negative control), FFD or TFD was also analyzed by solid phase binding. **c.** RAW264.7 cells were incubated with biotin-labelled TNF-α in the absence or presence of TNF antibody (positive control), FFD (10 μM) or TFD (1 μM) for overnight, then cells were analyzed by flow cytometry.

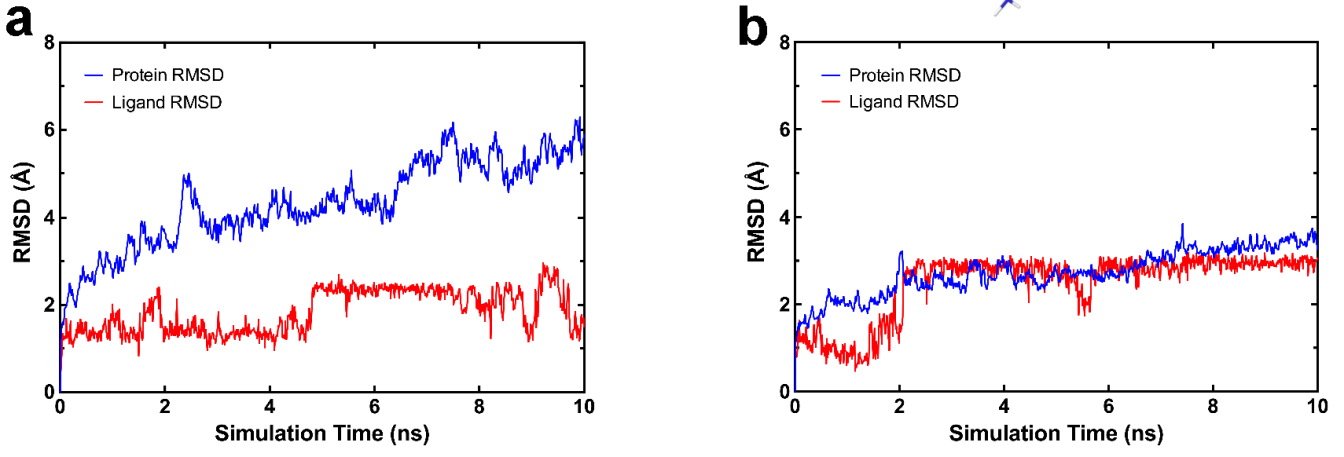

**Fig. S13. a.** RMSD trajectory of cPLA2 and Fexofenadine in the cPLA2-Fexofenadine complex over the 10 ns MD simulation. **b.** RMSD trajectory of cPLA2 and Terfenadine in the cPLA2-terfenadine complex over the 10 ns MD simulation.

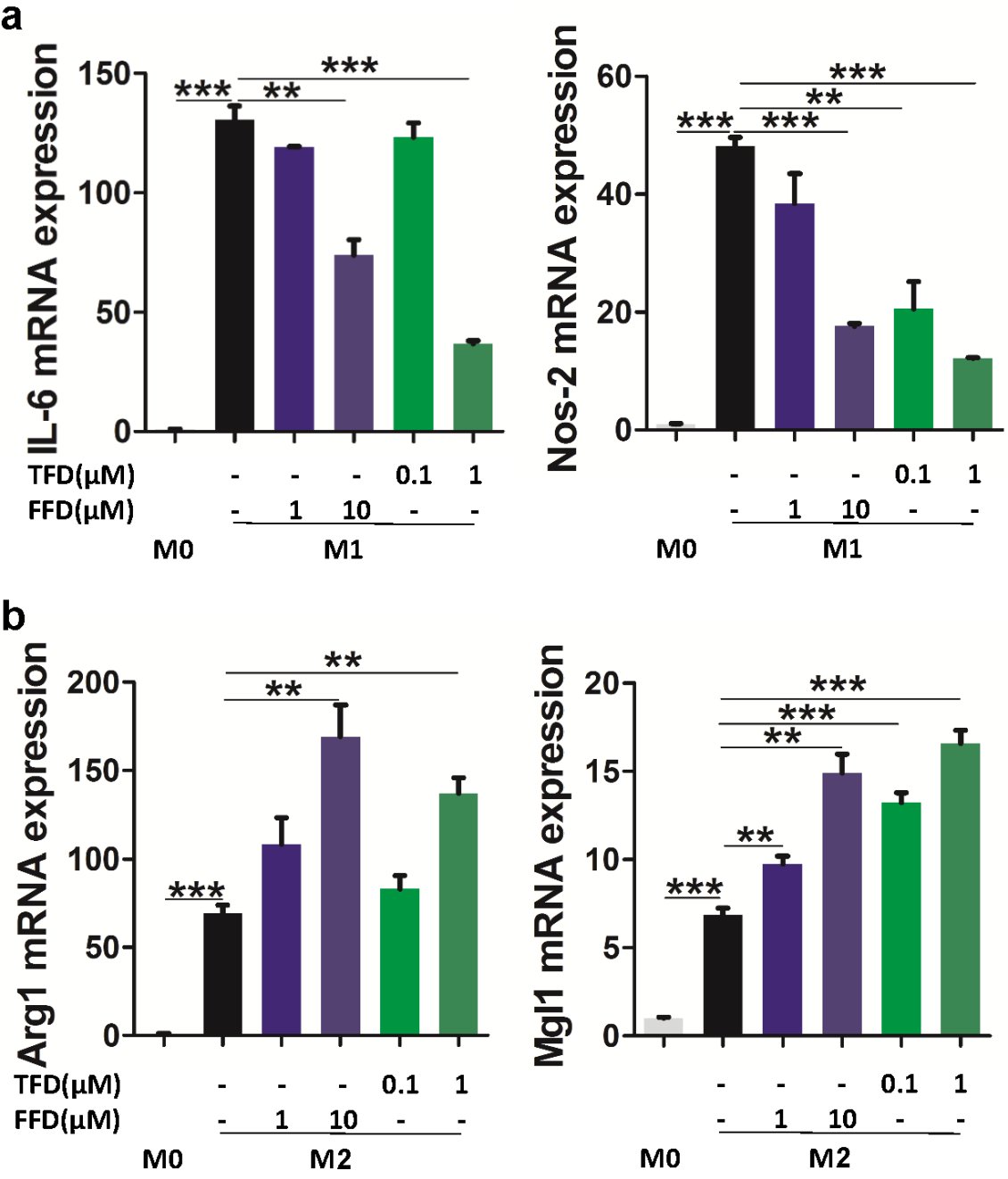

**Fig. S14. The effect of Fexofenadine (FFD) and Terfenadine (TFD) on macrophage polarization.** BMDMs were treated with M-CSF, IFN-γ (25 ng/ml) and LPS (250 ng/ml) or IL-4 (20 ng/ml) for type 1 macrophages (M1) or type 2 macrophages (M2) respectively. Various amounts of FFD or TFD, as indicated, were added. **a.** qPCR analysis of *Il6*, *Nos2* mRNA expression in BMDMs polarized to M1 macrophages. **b.** qPCR analysis of *Arg1 or Mgl1* mRNA expression in BMDMs polarized to M2 macrophages. (** p<0.01, ***p<0.001)

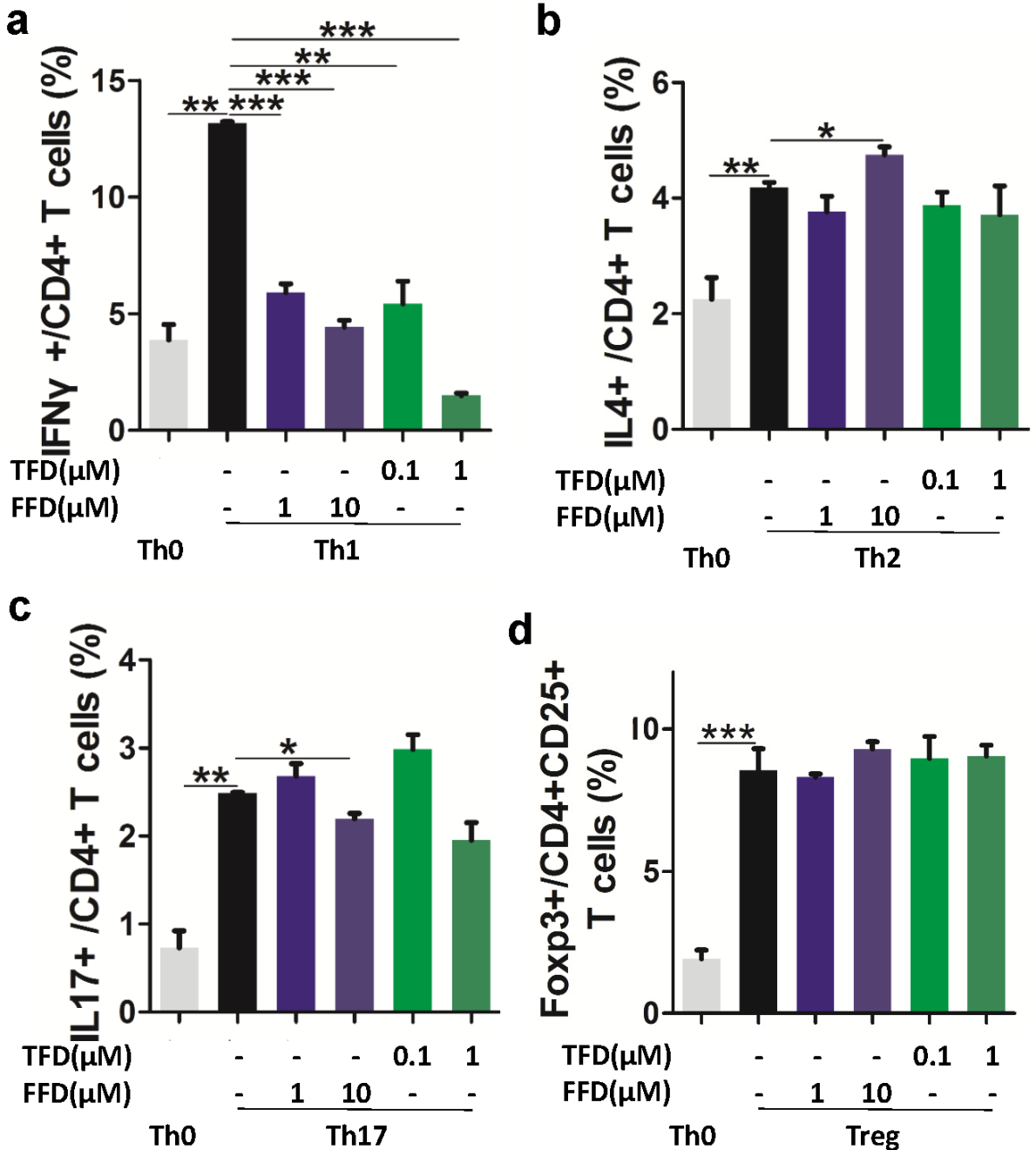

**Fig. S15. The effects of Fexofenadine (FFD) and Terfenadine (TFD) on T cell differentiation.** CD4+ T cells were isolated from spleen cells to determine the differential induction of T cell subtypes in the presence of various amounts of FFD or TFD, as indicated, for 4 days, flow cytometry was performed to examined the differentiation of Th1 (**a**), Th2 (**b**), Th17 (**c**) and Treg (**d**) cells. (* p<0.05, ** p<0.01, ***p<0.001)

**EXPERIMENTAL PROCEDURES**

**Screening FDA approved drugs *in vitro*.** NF-κB-bla THP-1 cell line (K1662, Invitrogen) was cultured in 96-well plates and FDA approved drugs (10μM, L1300, Selleckchem) were added and cultured overnight. Next day, cells were incubated with TNF-α (10ng/ml, PHC3015, Invitrogen) for 6 hours and LiveBLAzer™ FRET-B/G Loading Kit with CCF2-AM (K1095, Invitrogen) was used to detect the β-lactamase activity at wave lengths of excitation 409nm and of emissions 520nm or 470nm with SpectraMax i3x system (5025027A，Molecular Devices). This process was performed in triplicate. Subsequently, the drugs from first screening that could inhibit TNF-α induced NF-κB activity were undergone further screening by NF-κB luciferase assay. After transfected with plasmids of NF-κB luciferase reporter gene and Renilla for 8 hours, cells were treated with drugs (10μΜ) overnight, followed by TNF-α (10ng/ml) stimulation for 6 hours, then bioluminescence was measured by Dual-Luciferase® Reporter Assay kit (E1910, Promega). The data were analyzed by Graphpad.

**Screening and confirming the drugs' anti-TNF activity *in vivo*.** TNF-tg mice and NF-kB luc mice were purchased from Jackson Laboratory, and housed in Skirball Animal Facility of New York University Langone Medical Center. All animal experiments have been approved by Institutional Animal Care and Use Committee (IACUC) of New York University School of Medicine. We generated the TNF-tg/NF-kB-Luc mice by mating TNF-tg mice with NF-kB-Luc mice. After genotypes were confirmed by genotyping and bioluminescence via IVIS, the TNF-tg/NF-kB-Luc mice were orally treated daily for 7 days, with one of the 8 drugs selected in screen two. The luciferase activity was detected by IVIS 15 minutes after the injection of D-Luciferin (LUCK-1G, Gold biotechnology).

**Primary bone marrow derived macrophages (BMDMs) extraction.** BMDMs were isolated from C57BL/6 and TNF-tg mice. After mice were sacrificed by cervical dislocation, the tibia and femur were isolated and both ends of the bones were cut to open bone medullary cavity. Bone marrow cells were collected by centrifuge at 13000g for 90 seconds and seeded in 6-well plates. Prior to their use in experiments, M-CSF (10ng/ml, 576406, Biolegend) was added for 6 days to the BMDM culture medium.

**qRT-PCR.** Total RNA was extracted by RNeasy plus mini kit (74106, QIAGEN) and cDNA was synthesized using SuperScript® Reverse Transcriptase (M314C, promega Corporation). SYBR® Green PCR Master Mix (4309155, Applied Biosystems) was used to perform quantative real-time PCR (qRT-PCR) and the reaction was performed on a StepOnePlus^TM^ real-time PCR Systems (Applied Biosystems). The mRNA expression level of target gene was calculated by ΔΔCT and fold changes of mRNA levels were normalized to GAPDH.

**ELISA.** Cytokine levels of IL-1β and IL-6 in cell cultural supernatants or sera from mouse models were detected by sandwich ELISA according to product specifications in ELISA kit (IL-1β 88-7013, Invitrogen; IL-6: 88706476, Invitrogen). Cell cultural supernatants were collected when cells were treated with drugs for 48 hours. Sera were separated from whole blood by centrifuge freshly collected blood from mice at 3000 rmp for 10 minutes.

**Western blot.** After protein samples were prepared, samples were separated by SDS-PAGE and transferred to a nitrocellilose (NC) membrane (162-0115，BIO-RAD) using a wet transfer system. Membrane was blocked in 5% (w/v) non-fat milk in TBST for half an hour at room temperature, followed by incubated with first antibody overnight at 4 degree and secondary antibody 1 hour at room temperature. The bands on the membrane were developed by chemiluminescent (ECL) substrate and visualized by GelDoc system.

**RNAseq and Transcription factors enrichment analysis.** BMDMs were incubated with Fexofenadine (10μM, S3208, Selleckchem) or Terfenadine (1μM, T9652, Sigma) with or without TNF-α (10ng/mL) for 24 hours. Total RNA was extracted by RNeasy Mini Kit (74106, Qiagen), and gene expression profiling analyzed by RNA-seq by NYU Genome Technology Center for RNA sequencing (Illumina HiSeq4000 Sequencing, HiSeq 4000 Single Read 50 Cycle Lane). In addition, TNF-α induced genes that were suppressed by FFD/TFD were used for Transcription factors enrichment analysis with TFactS^1^.

**Osteoclastogenesis.** BMDMs were obtained as described above and the resultant preosteoclasts were cultured in medium supplemented with 10 ng/ml TNF-α and 100 ng/ml RANKL in the absence or presence of Fexofenadine or Terfenadine for 4 days. The cells were fixed with formalin and stained for tartrate-resistant acid phosphatase (TRAP) with a TRAP solution containing 100 mM sodium acetate buffer (pH 5.0), 50 mM sodium tartrate, 0.1 mg/ml sodium naphtol AS-MX phosphate, 0.6 mg/ml Fast Violet LB, and 0.1% Triton X-100. TRAP-positive cells appeared dark red and TRAP-positive multinucleated cells containing more than three nuclei (TRAP-MNCs) were visualized using light microscopy.

**Nuclear translocation and DNA binding activity of NF-κB.** BMDMs were treated with or without Fexofenadine (10μΜ) or Terfenadine (1μΜ) overnight followed by 6 hour incubation with TNF-α (10ng/mL). Immunofluorescence was performed to test the location of p65 (4764s, cell Signaling). The cytoplasmic and nuclear protein fractions were extracted with a hypotonic solution and hypertonic solution. p65, Lamin B and GAPDH (2118, Cell Signaling) were detected in cellular fractions by Western blot. The total proteins were extracted for analysis of p65 DNA binding activity by TransAM® NFκB p65 ELISA kit (40096, Active motif).

**hTNF-tg mouse model**

Human TNFα gene was transfected to C57BL/6 background mouse, named human TNF transgenic (TNF-tg) mouse, which highly expresses TNFα and spontaneously develops arthritis^2^. For prevention assessment, oral treatments of Fexofenadine (10mg/kg), Terfenadine (50 mg/kg), MTX (2 mg/kg, serving as a positive control) were started at 8-week age for a total of 13 weeks. Treatments were stopped at 17-weeks-time point and resumed at 19-weeks-time point to observe the response of inflammatory arthritis progression to Fexofenadine or Terfenadine. For treatment assessment, treatments of Fexofenadine (0.4, 2, 10 mg/kg), Terfenadine (2, 10, 50 mg/kg), MTX (2 mg/kg) were started to orally deliver when the average swelling score reached approximately 8 points for a total of 8 weeks. Six mice in each group. Swelling scores were weekly assessed based on the scoring system offered by the company. The swelling scores were added by the scores from digits, paws, wrists and ankles and the highest score for each mouse was 24. The score system is: 20 digits (0=normal, 0.2= one or more swollen joints), 4 paws (0=normal, 1= noticeable swollen, 2=severe swollen), 2 wrists (0=normal, 1= noticeable swollen, 2=severe swollen), and 2 ankles (0=normal, 2= noticeable swollen, 4=severe swollen). At the end of treatment and observation, sacrificed the mice and collected sera, spleens and ankles.

**Collagen induced arthritis model.** Eight-week old male DBA/1J mice were bought from Jackson Lab. Emulsion of complete Freund's adjuvant (7001, Chondrex) and chicken type II collagen (20012, Chondrex) was intradermally injected at the site 1.5-2 cm distance from the tail base. For prevention assessment, oral treatments of Fexofenadine (10mg/kg), Terfenadine (50mg/kg), MTX (2 mg/kg, serving as a positive control) were started at 18th day after immunization for a total of 48 days. For treatment assessment, treatments of Fexofenadine (0.4, 2, 10mg/kg), Terfenadine (2, 10, 50mg/kg), MTX (2 mg/kg) were started to orally deliver when the average clinic score reached approximately 5 points for a total of 24 days. Eight mice in each group. Clinic score was recorded every other day based on the following sore system^3^: 0=normal, 1=mild swelling involving ankle, wrist, or one digit, 2= mild swelling involving entire paw or more than two digits, 3=moderate swelling from the ankle/wrist to entire foot/paw and all digits, 4= severe swelling at the whole ankle/wrist, foot/paw and digits or ankylosing deformity. After the mice were sacrificed, sera and joints were collected for further detection.

**DSS induced colitis model.** Dextran sulfate sodium (DSS)-induced colitis model was established by feeding the 8-weeks C57BL/6 mice with drinking water containing 3% DSS for 5 days and followed by normal drinking water for 3 days. 3 days before intervention, oral treatments of Fexofenadine (0.4, 2, 10 mg/kg), Terfenadine (2, 10, 50 mg/kg), 5-ASA (50 mg/kg) were started until the mice were sacrificed, 6 mice in each group. Body weight, stool consistency and rectal bleeding were recorded every day based on the scoring system as below^4^. Weight loss: 0= less than 1%, 1= between 5% and 10%, 2= between 10% and 15%, 3= between 15% and 20%, 4= over 20%. Stool consistence: 0=normal, 2=loose stool, 4=diarrhea. Rectal bleeding: 0=negative, 2=blood trace, 4=gross blood. Colon length was measured, and colons and the sera were collected for further detection.

**TNBS induced colitis model.** Before establishing trinitro-benzene-sulfonic acid (TNBS) induced colitis model, 150μL 1% TNBS was absorbed by the skin on the back between the front legs in 8-week old C57BL/6 mice to pre-sensitize. A week later, 100μL 2.5% TNBS was delivered through anus. Three days before molding, the oral treatments of Fexofenadine (0.4, 2, 10 mg/kg), Terfenadine (2, 10, 50 mg/kg), 5-aminosalicylic acid (5-ASA, 50 mg/kg, serving as a positive control) were started and lasted for 5 days, 6 mice in each group. Body weights were recorded every day. After the mice were sacrificed, colon length was measured, and colons and the sera were collected for further detection.

**Histology and analysis.** Tissues were fixed in 4% formaldehyde, decalcified in 10% EDTA and embedded in paraffin. Serial 5µm sections were cut and stained with hematoxylin and eosin (H&E). Images were obtained by digital microscope (Axio Scope A.1, Carl Zeiss, LLC) and scored by two independent observers. The scoring systems for ankles were consisted of inflammation and pannus formation and articular cartilage damage: Inflammation was scored as follows: 0, Normal, 1, local inflammatory infiltration and 2, marked infiltration with lymphoid aggregates and edema. Pannus formation and articular cartilage damage were scored as follows: 0, Normal, 1, synovial proliferation adjacent to cartilage but no articular cartilage damage and 2, synovial proliferation and articular cartilage damage. For histology analysis of colitis, the scores were the combination of inflammatory cell infiltration and intestinal wall structure integrity, the higher scores indicated more serious inflammation^5^. Scores of inflammatory cell infiltration were: 0=normal, 1=inflammatory cell only infiltrated to mucosa, 2= inflammatory cell reached to mucosa and sub-mucosa, 3=inflammatory cell was found in the whole intestinal wall. Scores of intestinal wall structure integrity were: 0=normal, 1= inflammatory cell was locally infiltrated, 2= focally formed ulceration, 3=extensively formed ulceration with or without granulation tissue or pseudo-polyps.

**Micro-CT.** Ankle tissues were fixed by 4% formaldehyde and stored in 70% ethanol. After fixation, tissues were scanned, at a resolution of 10.5 μm, by Scanco vivaCT40 cone-beam scanner (SCANCO Medical, Switzerland) with 55 kVp source and 145 μAmp current. The scanned images from each group were evaluated at the same thresholds to allow 3-dimensional structural reconstruction of each sample.

**TRAP staining.** For paraffin-embedded slides, the sections were firstly deparaffined in a xylene and ethanol gradient, and then stained with TRAP staining solution mix for 60 minutes at 37°C and counterstained with methylene blue for 5 minutes, followed by dehydrating through graded ethanol and xylene. For live cultured cells, cells were firstly fixed by 10% formaldehyde for 10 minutes at room temperature, then were performed staining. The images was taken by light microscope (Axio Scope A.1, Carl Zeiss, LLC). The osteoclasts were stained with red violet in the green background.

**Safranin O staining.** After the paraffin-embedded knee and ankle sections were deparaffined in a xylene and ethanol gradient, sections were stained with with 2% hematoxylin (23412, MilliporeSigma ) for 5 min, 1.0% Safranin O (S8884, Sigma-Aldrich) for 60 min, and counterstained with 0.02% Fast Green (F7258, Sigma-Aldrich) for 1 minute. Stained slides were dehydrated, cover slipped and photographed.

**DARTS assay.** Drug affinity responsive target stability (DARTS) assay was performed based on previously reported methods^6^. In brief, cells lysate was extracted by M-PER™ Mammalian Protein Extraction Reagent (78501, Thermo Fisher), briefly centrifuged, and mixed with drugs or DMSO for 1 hour on a rotator. Pronase (P5147, Sigma-Aldrich) was added in the mixture for 15 minutes at room temperature. Digestion was stopped by 10 minute incubation with protease inhibitor cocktail on ice. Samples were boiled in SDS loading buffer for SDS-PAGE and Western blot.

**CETSA assay.** RAW264.7 cells were treated with FFD (10 μM)/TFD (1 μM) for 1 hour in a 37°C incubator with 5% CO_2_. Cells were harvested and divided equally into different tubes, after which the cells were heated at different temperatures for 3 minutes. After lysis via 3 repeated freeze-thaw cycles, samples underwent centrifugation at 20000g and supernatants were collected for analysis. For the isothermal dose response, after a series of ten-fold change concentrations ranged from 0.001 μM to 100 μM of FFD/TFD were used to treat cells and no variation in temperature was introduced. The remaining steps are the same as previously reported^7^.

**Site-directed mutagenesis.** PLA2G4A cDNA from Genscript was used as a template and Ser-505 was mutated to Ala using Q5® Site-Directed Mutagenesis Kit in accordance with the manufacturer’s instructions (E0554, New England Biolabs). To specific, taking PLA2G4A cDNA clone from Genscript as a template, the base of T was mutated to G in the amino acid Ser-505 (TCT) which changed to Ala505 (GCT).

**Construction of plasmids.** The construction of cPLA2 mutant plasmids was based on serial C-terminal and N-terminal deletion of the amino acid sequences associated with functional structure. Each plasmid carried a flag tag. The constructions were: full length cPLA2 (1-750), cPLA2 (126-750), cPLA2 (406-750), cPLA2 (1-479), and cPLA2 (1-144).

**cPLA2 activity assay.** Raw264.7 cells were transfected with an expression plasmid that encoding cPLA2. 24 hours later, the transfected cells were treated with TNF-α (10 ng/ml) and ATK (1 μM), or TFD (0.1 μM or 1 μM), or FFD (1 μM or 10 μM) overnight. Then the cells were collected and lysed and the cell lysate was used to do the measurement of cPLA2 activity using ELISA kit (765021, Cayman). Briefly, the protein concentration of the cell lysate was measurement by BCA, each sample with the same amount protein was incubated with cPLA2 substrate (Arachidonoyl Thio-PC) at room temperature. One hour later, DTNB / EGTA was added to stop the enzyme catalysis, and then the absorbance was read at 414 nm using a plate reader.

**Knock down of H1R1 by siRNA.** BMDMs were transfected with siH1R1, or scrambled control siRNA (scRNAi) using lipofectamine 2000 (11668-019, Invitrogen) according to manual specification. After 24 hours, supernatants and lysate were collected to detect cytokine secretion and protein expression, respectively.

**Knock out of cPAL2 by CRISPR-Cas9.** Knockout cells were generated in accordance with a previously published protocol^8^. gRNA was inserted it into the lentiCRISPR v2 vector. Packaging the inserted vector into lentivirus was completed by co-transfecting the lentiCRISPR v2, VSVG and ΔR6.7 in 293T cells. After packaging, we transfected the virus into RAW264.7 cells over 8 hours. Puromycin (2 ug/ml) was used to select the stably transfected RAW264.7 cells

**Arachidonic acid detection.** BMDMs were treated with TNF-α (10 ng/ml) with or without Fexofenadine/Terfenadine/ATK, the cultured medium was collected and analyzed using an arachidonic acid ELISA kit (NBP2-66372, Novus Biologicals). The result was detected at 450nm by automatic plate reader. ATK was implemented as a positive control.

**Solid phase binding.** To test whether Fexofenadine/Terfenadine affect the binding between TNF-α and TNFR1, solid phase binding assay was performed according to previously described method^9^. Briefly, a 96-well ELISA plate was coated with 100 μL of 0.5 ng/μL TNFR1 at 4℃ overnight. The plate was washed with PBST 5 times, and 300 μL blocking buffer was added to each well followed by incubation at room temperature for 1 hour. After discarding the blocking buffer, 50 μL buffer containing drugs at concentrations ranging from 0.1 nM to 10^5^ nM were added to the plate for 1 hour at room temperature. Three wells incubated with BSA served as negative controls and three wells containing TNF-α were set as positive controls. Without washing, 50 μL buffer containing 10 ng biotin-labeled TNF-α was added to each well, and incubated at room temperature for 2 hours. The plate was washed and 100μL buffer containing streptavidin-HPR was added and incubated at room temperature for 30 minutes. After a final wash, 100μL TMB buffer was added to each well and the reaction was stopped when the positive control group turned blue. The plate was read by automatic plate reader at 450nm.

**Flow cytometry.** To test whether Fexofenadine/Terfenadine affected the binding between TNF-α and TNFRs expressed on cell surface, flow cytometry was performed. Raw 264.7 cells were seeded in a 12-well plate. After adding DMSO (control group), Fexofenadine (10 μM) or Terfenadine (1 μM) overnight, cells were stained according to manual specifications (NFTA0，R&D Systems). Then the samples were analyzed at NYU core facilities using a FACSCalibur cell analyzer with CellQuest software.

**Spleen CD4^+^T cell differentiation.** Inducing spleen naïve CD4^+^T cell differentiation into T cell subsets was performed based on previously published methods^10^^,^^11^. Briefly, after preparing the naïve CD4^+^T cells, the following cytokines were added to induce spleen naïve CD4^+^T cell differentiation into T cell subsets: IL-2 (20 ng/mL), IL-12 (15 ng/mL), and anti-IL4 (5 μg/mL) for Th1; IL-2 (20 ng/mL), IL-4 (10 ng/mL), and anti-IFN γ (5 μg/mL) for Th2; IL-6 (20 ng/mL), TGFβ (3 ng/mL), anti-IFN γ (5 μg/mL), and anti-IL4 (5 μg/mL) for Th17; IL-2 (20 ng/mL), TGFβ (15 ng/mL), anti-IFN γ (5 μg/mL), and anti-IL4 (5 μg/mL) for Treg. Fexofenadine (1 or 10 μM) or Terfenadine (0.1 or 1 μM) were added and cells were cultured for 4 days without changing the medium.

Before collecting cells for fluorescence staining, 1 μL Golgi stop from Fixation/ Permeabilization Solution Kit with BD GolgiPlug™ (555028，BD Biosciences) was added to each well and incubated the plate for 4 hours. Cells were stained according to the kit specifications. For each T cell subtype, the combination of fluorescence dyes was as follows: FITC-CD4 (M1004502, Sungene) and Percp-cy5.5-IFNγ (45-7311-82, eBioscience) for Th1; FITC-CD4 and PE-IL4 (554389, BD pharmingen) for Th2; FITC-CD4 and PE-IL-17A (12-7177-81, eBioscience) for Th17; FITC-CD4, PE-CD25 (**102008**, Biolegend) and Alex-Fluo647-FoxP3 (51-5773-80, eBioscience) for Treg. Then the samples were analyzed at NYU core facilities using a FACSCalibur cell analyzer with CellQuest software.

**Macrophage polarization.** To test the influence of Fexofenadine/Terfenadine on macrophage polarization, subtype marker expressions were tested by qRT-PCR. Nos2 and IL-6 were used to indicate M1 macrophage while Arg1 and Mgl1 were used to indicate M2 macrophage. qRT-PCR primer sequences of the markers were: Arg1 5’-3’: F-TGC CAA AGA CAT CGT GTA CAT TG and R-CTT CCC AGC AGG TAG CTG AAG. Mgl1 5’-3’: F-CAG ATC CGT ATC TGT CTG GAT C and R-AGG TGG GTC CAA GAG AGG ATG. Nos2 5’-3’: F-TGT TAG AGA CAC TTC TGA GGC TC and R- ACT TTG GAT GGA TTT GAC TTT GAA G. IL-6 5’-3’: F-TTC CAT CCA GTT GCC TTC TTG and R: AGG TCT GTT GGG AGT GGT ATC. INFγ (25 ng/mL) and LPS (250 ng/mL) were added to BMDMs to induce M1 polarization while IL-4 (20 ng/mL) was used to induce M2 polarization, with or without adding FFD (1 or 10 μM) or TFD (0.1 or 1 μM), and cells were cultured for 24 hours. mRNA were extracted by RNA Mini kit (74106，QIAGEN).

**Protein Modeling and Preparation.** The crystal structure of human cPLA2 (PDB ID: 1CJY, 2.5Å) is available, however, there are several missing regions (residues 1 to 12, 406 to 414, 433 to 459, and 499 to 538) in this crystallized protein that were not resolved in the X-ray diffraction^12^. The missing structural regions were modeled into chain A of 1CJY using Modeller v9.20 (Andrej Sali Lab, UCSF, CA, USA, 2018). The generated homology model of 1CJY was further equilibrated and refined using 50 ns of all atom MD simulations by Desmond v5.6. The refined protein structure was then prepared using Protein Preparation Wizard implemented in Maestro v11.1 (Schrödinger, Inc., NY, USA, 2017) for docking simulation.

**Ligand Docking and Molecular Dynamics (MD) Simulation.** The 3D structures of Fexofenadine and its parent drug Terfenadine were built using Maestro v11.1 and energy minimized using the Macromodel v11.5 module. The ligands were then prepared with Ligprep v4.1 module to generate low-energy 3D structures. Since the cell-based study implicated that the impeded phosphorylation on residue Ser-505 may be a key mechanism involved in the inhibitory effect of fexofenadine on cPLA2, a docking grid (25 Ǻ) was generated by selecting Ser-505 as centroid. Flexible docking Fexofenadine and Terfenadine into cPLA2 was carried out using the XP (extra precision) mode by Glide v7.4 (Schrödinger, Inc., NY, USA, 2017). Taking the flexibility of receptor and ligand into consideration to get optimal binding simulation between ligand and receptor, induced-fit docking (IFD) by Glide v7.4 was performed. The ligand binding pose with the lowest predicted ligand binding free energy got from Glide XP docking process was subjected to IFD simulation following the default IFD protocol. The Glide Emodel value was ranked to identify the best docked pose among multiple conformations^13^. To validate the prediction of the binding, the docked cPLA2-fexofenadine complex and cPLA2-terfenadine complex with the best Glide Emodel values were subsequently subjected to short molecular dynamics simulation with Desmond MD system v5.6 (D. E. Shaw Research, NY, USA, 2018; Schrödinger, Inc., NY, USA, 2018). Predefined TIP3P water model was used and counter Na+/Cl- ions were added to balance the system charge. Periodic boundary conditions were set up and the buffer distance between box wall and the protein-ligand complex was set to be greater than 15 Å to avoid direct interaction of the complex with its own periodic image. After establishing the solvated system, MD simulation was carried out in the NPT ensemble using OPLS 2005 force field. The temperature and pressure were retained at 300 K and 1 atmospheric pressure. A short equilibration phase simulation was involved using default Desmond protocol, followed by running the 10,000 ps (10 ns) MD simulation for the equilibrated complex system. Schrödinger simulation interactions diagram (SID) was used to evaluate the interaction between ligand and protein, and the root mean square deviation (RMSD) of the ligand-receptor complex was calculated to evaluate if there are conformational changes of the protein or internal fluctuations of the ligand^14^. The RMSD was calculated for all frames in the simulation trajectory, with respect to the first frame as the reference frame. All calculations mentioned above were performed on a 6-core Xeon processor except MD jobs which were performed on a Nvidia GPU.

**Statistical analysis**

Data were reported as the means with standard errors. Comparisons among the treatment groups were performed by repeated measures in SPSS software (IBM, Armonk, NY, USA) or unpaired t-tests/one-way ANOVA in Graphpad software (GraphPad Software, San Diego, CA). P value < 0.05 was statistically significant at two-sides.
